# Ictal-Related Chirp as a Biomarker for Monitoring Seizure Progression

**DOI:** 10.1101/2024.10.29.620811

**Authors:** Nooshin Bahador, Frances Skinner, Liang Zhang, Milad Lankarany

**Affiliations:** Krembil Research Institute, University Health Network, Toronto, Canada; Department of Physiology, University of Toronto, Toronto, Canada; Department of Medicine (Neurology), Institute of Medical Science, University of Toronto, Toronto, Canada; Institute of Biomedical Engineering and Department of Physiology, University of Toronto, Toronto, Canada

## Abstract

Despite being prevalent, the causes, mechanisms, and progression of epilepsy—a chronic neurological disorder with unprovoked seizures—are not well understood, complicating drug development for treatment. This study used a comprehensive mouse epilepsy kindling model dataset to investigate frequency modulation (chirp) as a potential indicator of distinct states of epilepsy (early evoked discharge, late evoked discharge, spontaneous recurrent seizure, and drug state). Employing time-frequency ridge extraction, chirp identification, and statistical testing, our analyses revealed that chirp patterns occur in the majority of ictal discharges (>81.6%), persisting across evoked and spontaneous seizures. While the focus was on hippocampal recordings, chirps were also detected in the piriform peripheral cortex. Significant frequency and duration changes in chirp patterns during the transition from early to late evoked ictal events suggest their potential as the screening tool for seizure progression. Additionally, detailed analyses illuminate the impact of Lorazepam, a *GABA*_*A*_ enhancer, on chirp characteristics, providing insights into how increased inhibitory tone quantifiably influences excitatory-inhibitory balances during seizures.

## Introduction

Seizures are episodes of abrupt, brief bursts of electrical activity within the brain, resulting from imbalances between excitatory and inhibitory neurotransmitters, which can lower a neuron’s firing threshold. A seizure represents a singular event, whereas epilepsy denotes a neurological disorder marked by the occurrence of two or more unprovoked seizures. Various factors, including low oxygen, genetic predisposition, age, and medical conditions, can trigger seizures, leading to different types based on which brain regions are involved. Symptoms vary widely, manifesting as twitching, confusion, or sensory disturbances, and etc and understanding these manifestations aids in diagnosis and treatment (***Freeman et al., 1993***).

Epileptiform EEG activity is classified as ictal, signifying its occurrence during a seizure, postictal, indicating the period following a seizure, and inter-ictal, representing the intervals between seizures. The ictal seizure event involves states of initiation, maintenance or propagation and termination, and epilepsy research tends to focus on one of these phases (e.g. ***Miri et al. (2018***) and ***Rich et al. (2020***) consider initiation phases from experimental and computational perspectives, respectively). In general, evidence suggests that initiation is associated with increased inhibitory synchronization leading to subsequent sparse neuronal firing and an increase in extracellular potassium (see Figure 4 in ***de Curtis and Avoli (2016***)). The many interacting dynamics can produce ‘chirp-like’ patterns during the ictal electrographic discharges, where a chirp is a consistent increase or decrease in frequency within a very narrow band, whether linear or nonlinear (***Grinenko et al., 2018***).

Chirps are found in signals across a wide range of biological and physical phenomena, including bird songs, insect communication, and radar systems. In fields like optics and telecommunications, chirps can emerge when a pulse undergoes manipulation during its propagation through a medium with dispersion properties (***Frosz and Andersen, 2007***). Studies from human data have reported that chirp-like patterns are present in electrographic discharges from the epileptic brain (***Li et al., 2020***; ***Gnatkovsky et al., 2011***; ***Kurbatova et al., 2016***; ***Sen et al., 2007***; ***Niederhauser et al., 2003***; ***Schiff et al., 2000***; ***Feltane et al., 2013***; ***Gnatkovsky et al., 2019a***; ***Benedetto and Colella, 1995***) Analysis of seizure outcomes confirms that the fast activity chirp pattern is a consistent biomarker of the epileptogenic zone in a diverse group of patients undergoing stereoelectroencephalography (SEEG) (***Di Giacomo et al., 2024***). Although both clinical and experimental studies express similar observations on the existence of a chirp-like pattern in time-frequency representations of electro-graphic seizure data, the reported timing of chirp occurrences differs, with some indicating onset of ictal discharge (***Li et al., 2020***; ***Gnatkovsky et al., 2011***; ***Kurbatova et al., 2016***; ***Sen et al., 2007***; ***Niederhauser et al., 2003***; ***Schiff et al., 2000***; ***Feltane et al., 2013***; ***Gnatkovsky et al., 2019a***), others during transition from inter-ictal to ictal states (***Kurbatova et al., 2016***), and some even prior to ictal discharges (***Niederhauser et al., 2003***; ***Schiff et al., 2000***; ***Feltane et al., 2013***; ***Gnatkovsky et al., 2019a***; ***Benedetto and Colella, 1995***). The frequency dynamics of chirps also differ, including linear frequency decrease (***Feltane et al., 2013***), upward-downward trend (***Li et al., 2020***), and even harmonic structures (***Schiff et al., 2000***; ***Feltane et al., 2013***; ***Gnatkovsky et al., 2019a***; ***Benedetto and Colella, 1995***). This wide variation is difficult to reconcile as there is a limited amount of available data to analyze, making it challenging to observe any consistent and statistically significant results. For example, the report of spectral chirps in ***Schiff et al. (2000***) relied on observations of 35 chirps in 42 seizures from 6 patients with no quantitative assessment of the primary characteristics of these chirps, specifically in terms of their frequency, duration, and onset. The potential count of ictal discharges for a human subject is limited to a small number (fewer than four) over the span of one month in an optimal scenario (***Trevorrow, 2006***; ***Reiter and Andrews, 2000***). A study identified the chirp as a key fingerprint of the epileptogenic zone in human, based on 51 individual seizures, with three clinical seizures analyzed for each of the 17 patients (***Grinenko et al., 2017***).

Rodent animal models can exhibit multiple ictal discharges per day, so that over the course of several months, a substantial volume of data can be available for analyses. In this work, we take advantage of this to fully characterize chirps in ictal discharges from long-term recordings of a kindled mouse model (***Bin et al., 2017***; ***Liu et al., 2021***). From our analyses, we obtained a substantial (>500) quantity of chirps. This enabled us to continuously track the dynamics of chirp patterns over time. Thus, although this is not the first time ictal-related chirps have been observed, it is the first time that reliable characterization of ictal-related chirp patterns has been achieved. We found statistically significant differences in onset, duration, and median frequency of chirps over time and with a GABAergic enhancing drug (lorazepam), suggesting that chirp patterns could serve as a biomarker monitor for tracking seizure progression and charcterizing the impact of drug treatments.

## Results

We used long-term continuous rodent EEG recordings from kindled mice (see Methods) to detect chirps. These data included evoked states (early and late), spontaneous recurrent seizure (SRS) states (early and late), and a drug (lorazepam) state and recovery. We focused on hippocampal recordings, although recordings in other brain structure regions were also examined. To automatically identify chirps, we developed a method to reliably infer chirp-like patterns. Our method identified chirps in more than 80% of ictal events, resulting in more than 500 chirps for subsequent analyses. The analysis revealed distinctive spectro-temporal morphologies associated with chirps, and a consistent temporal evolution of chirp characteristics with statistically significant changes. These dynamic changes in chirp characteristics thus present an opportunity for understanding and monitoring the evolution of seizure events.

### Chirp detection and observations

We developed a novel method for chirp detection. It involves determination of onset and offset times of ictal discharge-related chirps and consists of six steps, using time-frequency analyses and thresholding. These steps are described and illustrated in detail in the Methods. This method allowed us to identify over 500 chirps from the data, representing 81.6% of all of the ictal events. While electrode locations in each animal may slightly vary, the fundamental feature of chirp remain more or less consistent. FIGURE 1(top plot) shows the evolving frequency pattern of a sample extracted chirp over time.

**Figure 1.**
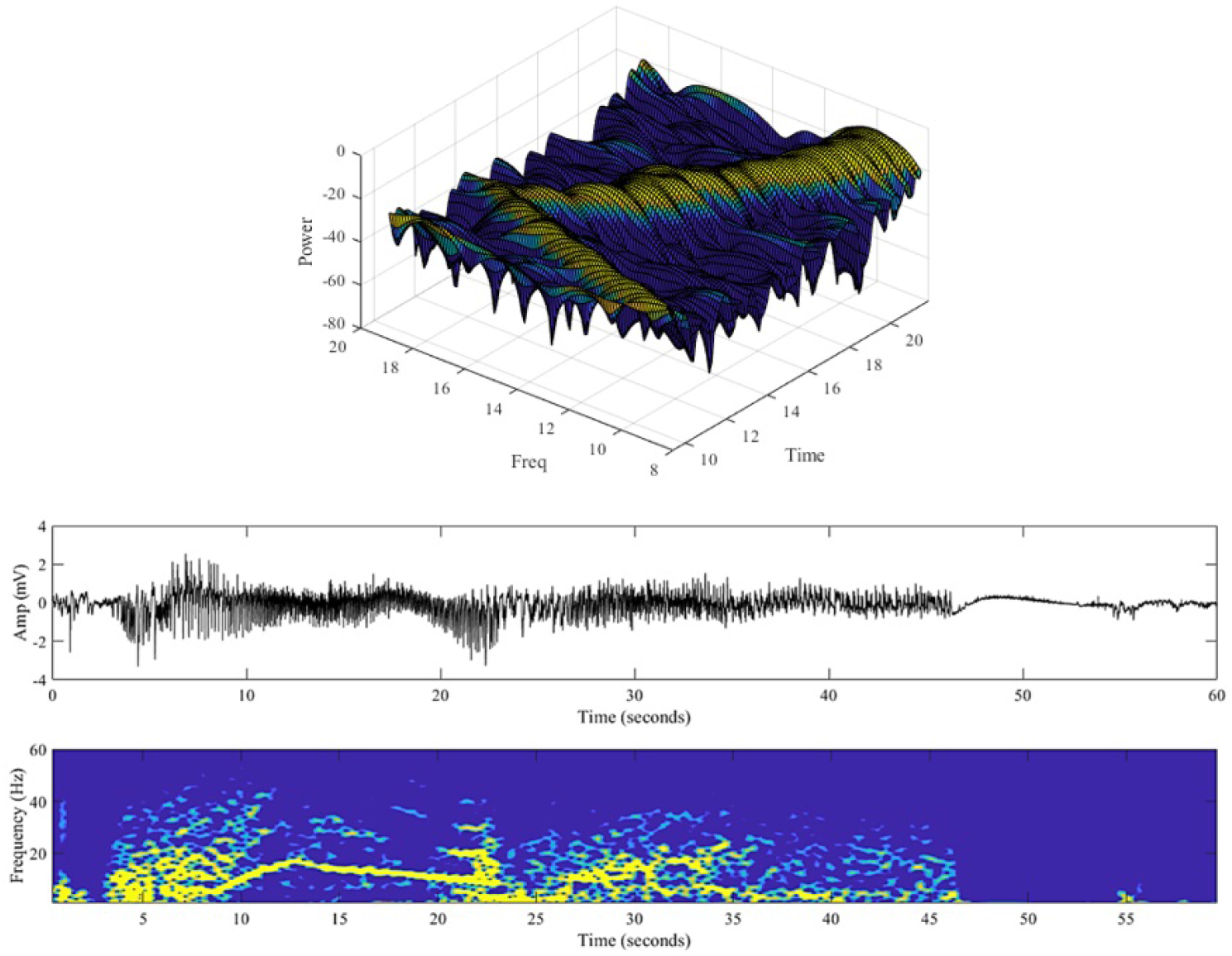
An example of a chirp pattern. 3D Visualization of a sample ictal discharge chirp with asymmetric peak-structured frequency variation over time is shown in the top plot. The accompanying recording with its 2D time-frequency representation is shown in the lower plot. Data from mouse Cage#7. **Figure 1—figure supplement 1. Chirps exhibiting harmonic relationships**. **Figure 1—figure supplement 2. Chirp patterns in two different brain regions**. **Figure 1—figure supplement 3. Spectro-temporal morphology of ictal chirps**.

Within certain ictal discharges, it was observed that the frequencies of individual chirps exhibited a harmonic relationship. See *Supplemental Figure 1* of FIGURE 1. The harmonically related chirps appear to originate within the same structure since otherwise, if they originated from different structural regions, there would be some noticeable delay. Chirp patterns were detected in brain structures other than the hippocampus. Specifically, we found comparable chirp patterns in the piriform cortex recordings that were simultaneously obtained with hippocampal recordings in the same animal. An example is shown in *Supplemental Figure 2* of FIGURE 1 for a SRS event. The data analysis findings showed a progressive evolution of chirp activities, transitioning from its initial formation to the emergence of a fully developed pattern over time. Furthermore, there were observations of recurring, distinct chirp patterns occurring in a cyclic manner over time. In *Supplemental Figure 3* of FIGURE 1, different spectro-temporal patterns of ictal chirps are presented.

Based on all of these observations, we believe that chirps are an inherent characteristic of ictal discharges regardless of being evoked or spontaneous. Thus, chirp activity can be viewed as a representative feature of a large local circuitry, as it was observed in all studied animals.

### Chirps as biomarkers

Considering the observed richness in chirp characteristics, several chirp features were automatically extracted with the intention of proposing their use as biomarkers. The automated characterization of ictal-related chirps involved analyzing chirp features of duration, median frequency, and onset (distance from ictal discharge onset), to better understand the temporal and spectral characteristics. FIGURE 2 shows an example of a spectrogram with its extracted chirp features.

**Figure 2.**
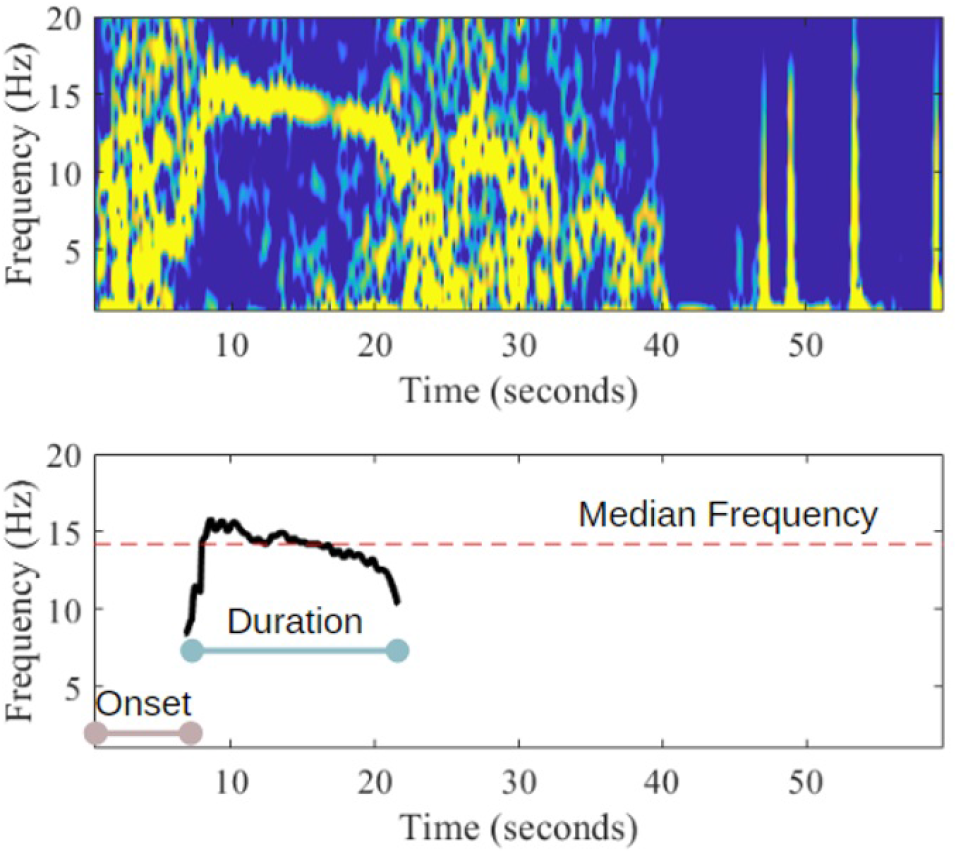
Illustration of chirp features. Spectrogram and corresponding extracted ictal-related chirp, with key features highlighted, including the chirp’s duration, median frequency, and chirp onset (distance from ictal discharge onset).

We found that chirp features changed with seizure progression over time. In FIGURE 3, we show the temporal evolution of mean values and their corresponding standard errors for characteristics of chirp-like patterns throughout different stages of epilepsy ictal events and a drug application. These states span from early evoked, late evoked, SRS, later SRS, and a drug state and recovery. The later SRS state is the ‘before drug’ state, but as the drug administration was later, it also allowed us to consider the evolution of SRS over time. The drug used is lorazepam, which is an anti-seizure medication. While it has strong peripheral action and is well-established as an effective agent for relaxing peripheral muscles and mitigating convulsive behavior (***Kienitz et al., 2022***), it may or may not suppress ictal discharges.

**Figure 3.**
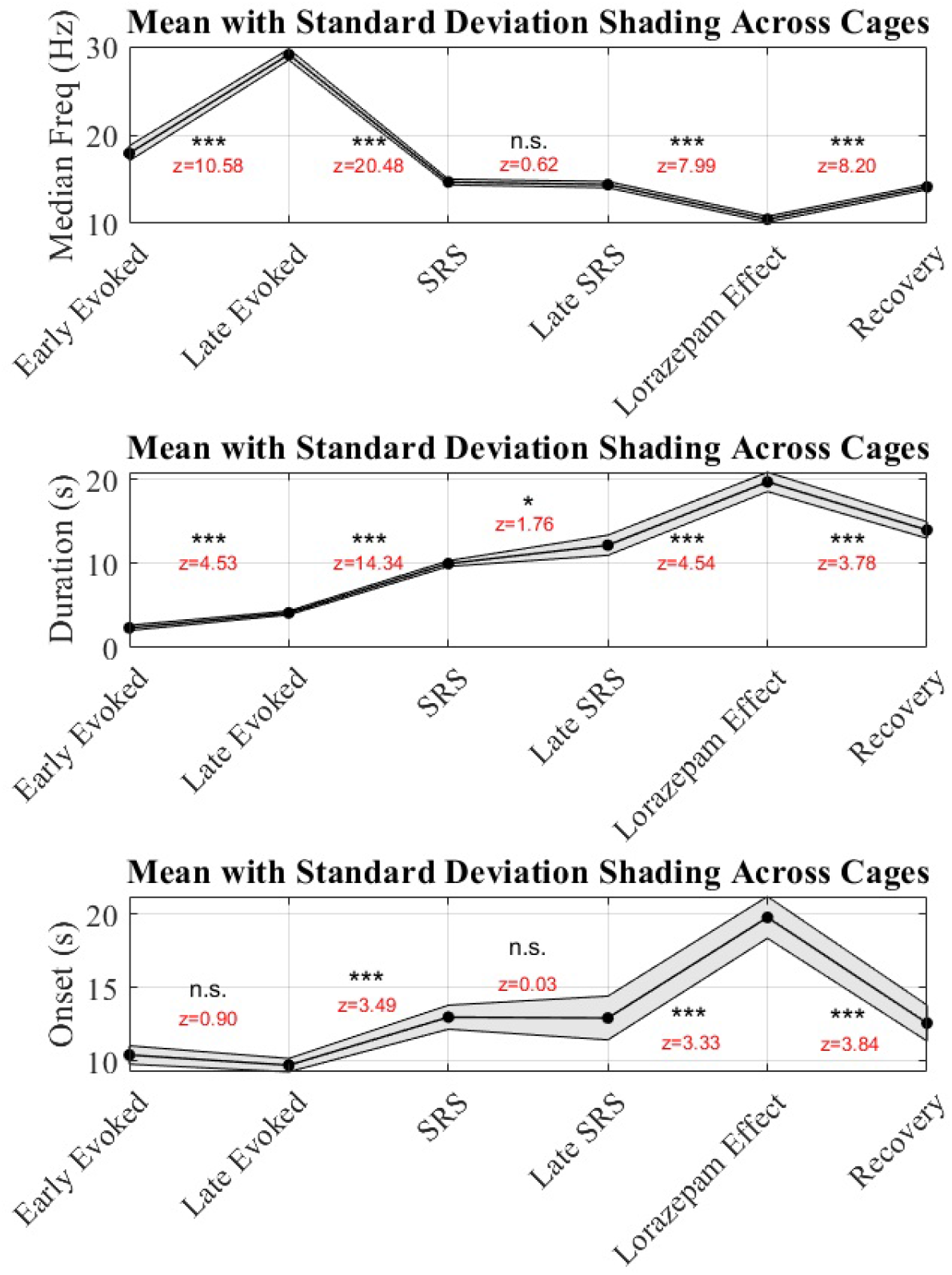
Evolution of chirp features. As epilepsy progresses from the initial evoked state to a drug state, significant changes in mean values and condence intervals are observed for chirp-like pattern characteristics. These mean values and their associated standard errors were obtained by conducting pairwise comparisons using a multiple comparison test to determine the states with statistically significant differences in means. The z-score was used as a measure to quantify the difference between two means relative to their variability. The calculated z-scores for each pair were classified into significance levels as follows: a z-score exceeding 2.58 provides strong evidence against the null hypothesis (*p* < 0.01), denoted by ***; a z-score above 1.96 indicates moderate evidence (*p* < 0.05), marked with **; and a z-score greater than 1.64 suggests some evidence (*p* < 0.10), indicated by *. If none of these thresholds are met, the result is labeled as “not significant” (n.s.). **Figure 3—figure supplement 1. Variation in chirp duration (s) across different animals over time: Early Evoked, Late Evoked, SRS, Late SRS, Drug, and Recovery**. **Figure 3—figure supplement 2. Statistical difference in chirp duration (s) across different states: Early Evoked, Late Evoked, SRS, Late SRS, Drug, and Recovery**. **Figure 3—figure supplement 3. Chirp duration of Pre-Drug, Drug, and Recovery states in different animals**.

We compared the median frequency in the six consecutive states statistically and found that each one significantly differed from the previous state, except for the comparison between SRS and Late SRS. Evoked ictal discharges occur artificially due to stimulation, which is possibly why their frequencies are higher compared to spontaneous ictal states (see median frequency plot in FIGURE 3). For chirp durations, there was a continual increase from the early evoked state to the drug state, and it was significantly different between early and late evoked, late evoked and SRS, and before and during drug states. We statistically compared the onset of chirp across six consecutive states and found significant differences between each state, except for the early evoked versus late evoked states and between SRS and late SRS. In late evoked discharges, the chirps initiate earlier than for the SRS state (see onset plot in FIGURE 3). Looking more closely at the changes in chirp median frequency from early to late evoked ictal discharges, we noted a consistent increase over time with a relatively stable rate of change. That is, with increased stimulation, the higher frequency band rises. This is shown in FIGURE 4. We also noted that chirps tended to occur just prior to termination for evoked discharges, whereas for spontaneous ictal events, chirps clearly occurred within the maintenance stage (middle). We show an example of this in *Supplementary Figure 1* of FIGURE 4. In other words, evoked discharges terminate abruptly following the chirp, while spontaneous ones do so less distinctly. It may be that chirps contribute to the termination of seizure in early stages. Overall, these findings underscore the dynamic alterations in chirp-like patterns associated with epilepsy progression and drug intervention, with statistical significance guiding the interpretation of each comparison.

**Figure 4.**
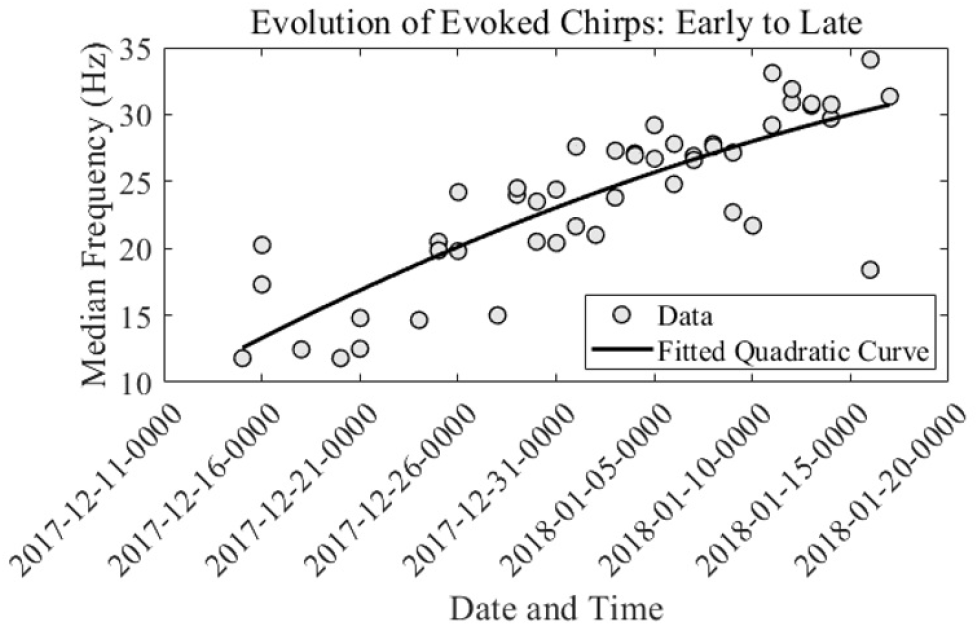
Evolution of evoked chirps. Changes in chirp median frequency from early to late evoked ictal discharges, showing a clear increase over time. Fit equation: *y* = −0.006169*x*^2^ + 9095*x* − 3.352 × 10^9^(*R*^2^ : 0.6811, Norm of residuals: 23.21) **Figure 4—figure supplement 1. Chirp occurrence differences**.

The series of t-tests (Two-Sample Assuming Unequal Variances) comparing chirp duration in different groups reveal several significant differences (The comparison involves animals from cage B #4, cage B #5, cage C #7, cage C #9, cage D #10, cage D #11, cage D #12, cage E #13, cage E #15, cage F #17, and cage F #19. see Tables of *Supplementary Table* 1 to *Supplementary Table* 7). The “Early Evoked” and “Late Evoked” groups show a significant difference with a *p*-value < 0.00001, indicating that the “Late Evoked” mean is significantly higher. Similarly, the “Late Evoked” and “SRS” groups differ markedly, with a *p*-value < 0.00001, showing that “SRS” has a significantly higher mean. The comparison between “SRS” and “Late SRS” also demonstrates a significant difference with a *p*-value < 0.00001. “Late SRS” and “Drug” show a significant difference as well, with a *p*-value < 0.01. Finally, “Drug” and “Recovery” are significantly different, with a *p*-value < 0.01, while “SRS” and “Recovery” also show a significant difference with a *p*-value < 0.00001. Comparisons between “Late SRS” and “Recovery” yield a marginally significant result with a *p*-value < 0.05.

Chirp duration in seconds of pre-drug, drug, and recovery states for various animals across different states was presented in *Supplementary Figure 3* of 3. The t-tests (Two-Sample Assuming Unequal Variances) assessed the duration differences between states: Pre-Drug, Drug, and Recovery. The results reveal that the mean duration in the Drug State (19.73 seconds) is significantly longer than in the Pre-Drug State (12.20 seconds), *p*-value < 0.001, indicating that the drug has a substantial effect on duration (see *Supplementary Table* 8). The mean duration in the Drug State (19.73 seconds) is also significantly longer than in the Recovery State (14.00 seconds). The two-tail *p*-value < 0.01 show a significant difference, suggesting the drug’s effect persists into the recovery period (see *Supplementary Table* 9). The mean duration in the Recovery State (14.00 seconds) is slightly longer than in the Pre-Drug State (12.20 seconds). However, the two-tail *p*-value < 0.1 indicate that while there is some evidence of a difference, it is not statistically significant at the typical 0.05 level (see *Supplementary Table* 10). So, the drug significantly increases the duration compared to both the Pre-Drug and Recovery States, but the difference between the Pre-Drug and Recovery States is less clear.

### Conceptual mechanisms underlying chirps

Our analysis results of distinct changes in chirp features indicate that there are mechanisms by which neurons synchronize to produce the narrow-band (NB) activity (i.e., chirps). We schematize a model mechanism conceptualization in FIGURE 5. Note that chirps occur during the maintenance phase of the ictal event, and we consider three other possible states. At the start of the seizure, there is broad-band (BB) activity which transitions (BB to NB) to chirps that terminate with BB activity.

**Figure 5.**
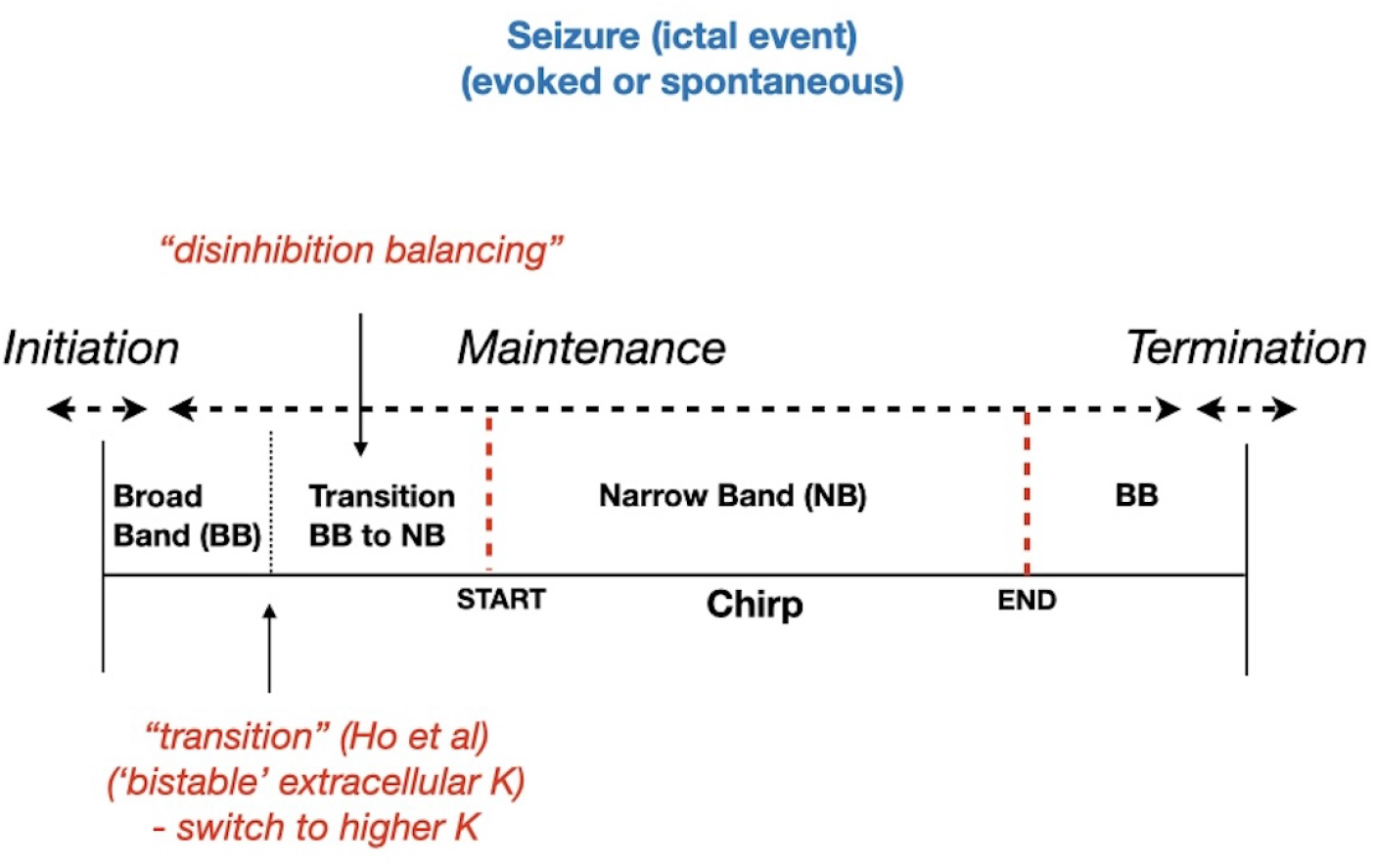
Model conceptualization schematic. Initiation, maintenance and termination phases are shown with chirps occurring the the maintenance phase. In our model conceptualization, we consider four states - a broadband (BB) irregular state following seizure initation, then the triggering of a narrower band state that eventually transitions to the narrowband (NB) chirp state. The last BB state may or may not be present before seizure termination (our observations indicate that for evoked states, this last BB state is non-existent).

Considering the chirp NB activity in the frequency domain, there must be significant synchronization in the underlying pyramidal cell population with some sufficient periodically occurring synaptic drive. That is, oscillating activity within excitatory-inhibitory (E-I) network circuits. We observed that chirps began earlier after multiple stimulations, comparing late versus early evoked states for chirp onset; however, this difference was not statistically significant. Initially (i.e., early evoked states), it presumably took longer to obtain chirps because at the early stimulation stage, neuronal recruitment was insufficient. For chirps to occur, a significant amount of wide-band activity is required. As more stimulations occur, the system somehow becomes more facilitative in generating this activity. Artificial stimulation induces evoked discharges, prompting the system to synchronize, allowing chirps to commence earlier. By enforcing synchronization, the process is expedited compared to the natural, spontaneous synchronization, which typically takes more time to occur.

To allow the NB activity to occur, a significant amount of BB activity is required. Although the exact location of chirps within an ictal discharge varies, it seems clear that chirps are associated with the maintenance stage of an ictal event. As previously summarized, seizure initiation is associated with increased inhibitory synchronization leading to subsequent sparse neuronal firing and an increase in extracellular potassium (***de Curtis and Avoli, 2016***). Moreover, we conceptualize that such synchronous inhibitory activity should evolve to suppress both principal (excitatory neurons) and inhibitory interneurons. This implies that initial inhibitory synchronization leads to sparse neuronal firing in principal neurons, reflected by a significant amount of wide-band activity, observed before (and after) narrow band activities(s) (chirp). Nevertheless, the same synchronous inhibition can be used to inhibit inhibitory interneurons, thus disinhibit principal neurons (with some delays).

Focusing on movement to the second state (transition from BB to NB, i.e., less BB) in the schematic of FIGURE 5, we invoke insights from previous E-I network models that also included dynamic extracellular potassium (***Y. Ho and Truccolo, 2016***). They were developed to understand different focal seizures observed in intracortical microelectrode recordings in humans (***Truccolo et al., 2011, 2014***). One of these seizure types is a ‘gamma seizure’ one, in which a narrow-band frequency arises, and there is a concomitant increase in extracellular potassium and asynchronous, heterogenous sparse firing of neurons. That global population rhythms could arise from a population of sparsely firing neurons has been described in classical theoretical studies (***Brunel, 2000***). The model mechanism underlying the narrow-band frequency development is due to interactions between the elevated extracellular potassium (due to abnormal potassium buffering) and the strength of synaptic inhibition, and is initiated by a brief perturbation (noise or stimulation in the model) due to bistable dynamics in high and low extracellular potassium states (***Y. Ho and Truccolo, 2016***). In our conceptualization, this transition along with disinhibition delays eventually leads to the start of a chirp. High inhibitory strength tends to decrease extracellular potassium in Ho’s model so that the duration of their gamma seizure is the result of a subtle balance between high inhibitory conductance and the accumulation of extracellular potassium. Further, earlier theoretical studies have shown that the population frequency in E-I networks decreases with longer (latency and decay) time constants of inhibition (see Figs 6/7 in ***Brunel and Wang (2003***)). The NB frequency aspect possible in ***Y. Ho and Truccolo (2016***)’s models arises from the previously ‘noisy’ wide-band ictal firing. We would consider this wide-band as the first state (BB) in our schematic of FIGURE 5 - such BB is seen in the recordings (e.g., see FIGURE 2). Then, a perturbation switches the system to higher extracellular potassium levels and our second state in the transition to narrow-band firing. In this conceptualization, the exact timing of chirp onset would vary depending on ‘noise and stimulation’ levels in the system (first state in schematic) along with delays associated with disinhibition (second state in schematic).

**Figure 6.**
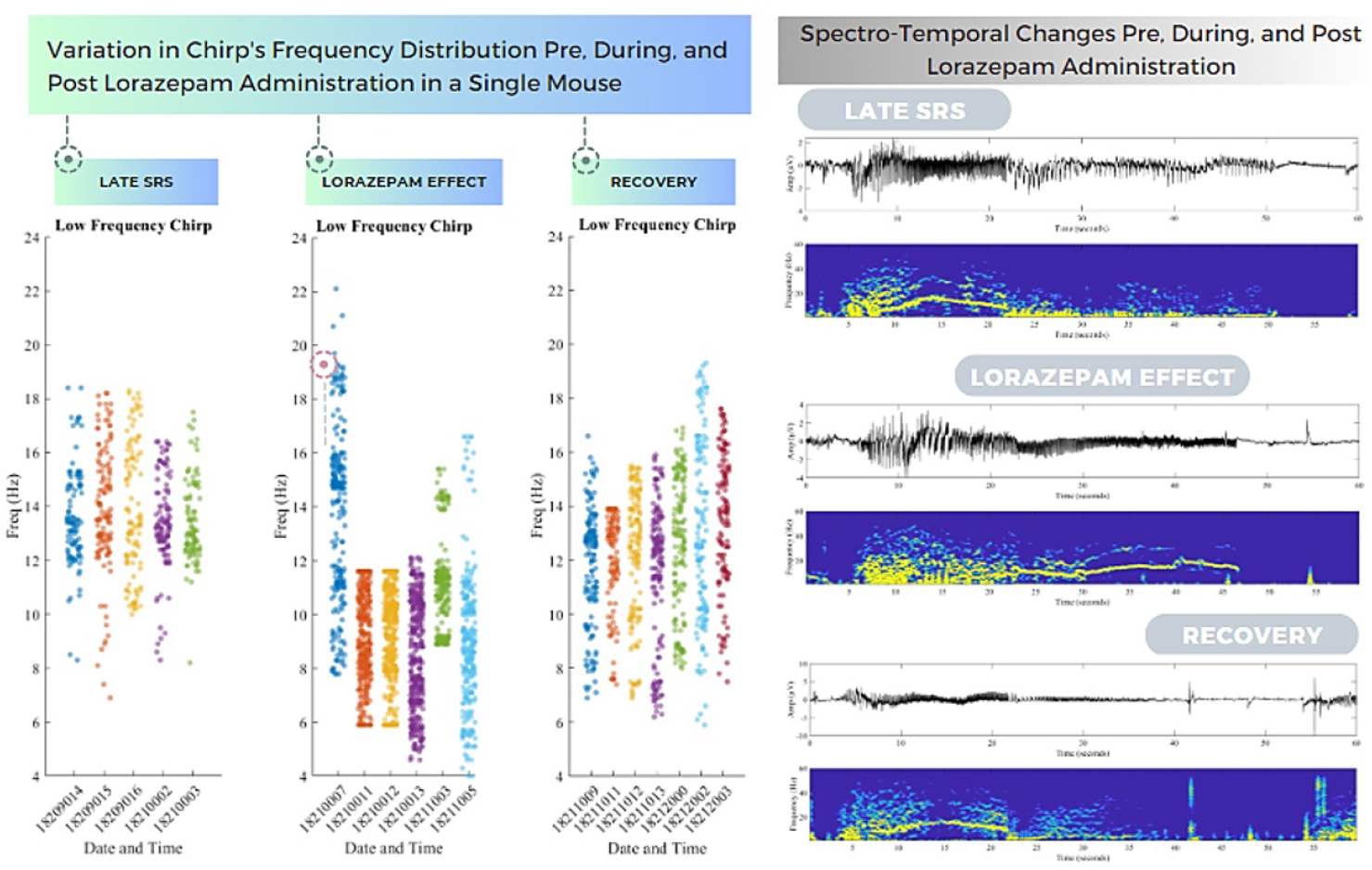
Chirp characteristics and drug application. (Left) Chirp frequencies are shown for the three states of: before lorazepam application (late SRS), during lorazepam, and after lorazepam (recovery). (Right) A single chirp is shown for the three different states to visualize the observed chirp characteristic changes.

Lorazepam is known to be a GABA_A_ (inhibitory) enhancer (***Chen et al., 2019***), and like other benzodiazepine GABA receptors, it increases inhibitory time constants (***Sanna et al., 2004***). Thus, based on our model mechanisms proposed above, increasing inhibition via lorazepam would affect E-I balances and distinctly affect chirp onset due to an increased time in the second state after a switch from BB. This is what we obtained in a statistically significant fashion (see FIGURE 3 onset plot). This can be more precisely seen for one of the kindled mice in FIGURE 6 (right). FIGURE 6 (left) shows that there is a clear decrease in chirp frequency and as time progresses after drug application, there is a gradual and upward shift in the chirp’s frequency that becomes noticeable during the extended recovery period. Following drug application, it is presumed that the first chirp still reflects the residual impact of the drug. However, as the recovery period lengthens, the drug’s influence gradually dissipates. Initially, the chirp is characterized by a confined frequency range, which then broadens as the recovery progresses. As the ictal discharge gradually recovers, continuous recording during the recovery phase ultimately leads to the chirp frequency back to levels that were present before drug injection. On the right of FIGURE 6, a single chirp from the three states of pre-drug (late SRS), lorazepam drug, and recovery is shown - a lengthening of chirp duration and chirp frequency decrease is clearly visible. Before the drug is given, chirp characteristics are considered as ‘baseline’ and used as a reference point to compare against the changes that occur after drug administration. When the drug is administered, it influences the chirp characteristics in various ways as shown. Monitoring the chirp characteristics after drug provides insights into how the seizure activity returns to its baseline state and whether there are any lasting effects from the drug exposure. That is, monitoring the chirp characteristics after drug allows one to gauge when the seizure activity returns to its baseline state and whether there are any lasting effects from the drug exposure.

Statistical analysis of data from all available animals that had lorazepam application revealed significant differences in the characteristics of chirp before, during, and after lorazepam administration. Specifically, lorazepam significantly extended the duration, reduced the median frequency, and prolonged the onset time of chirps. (see FIGURE 7).

**Figure 7.**
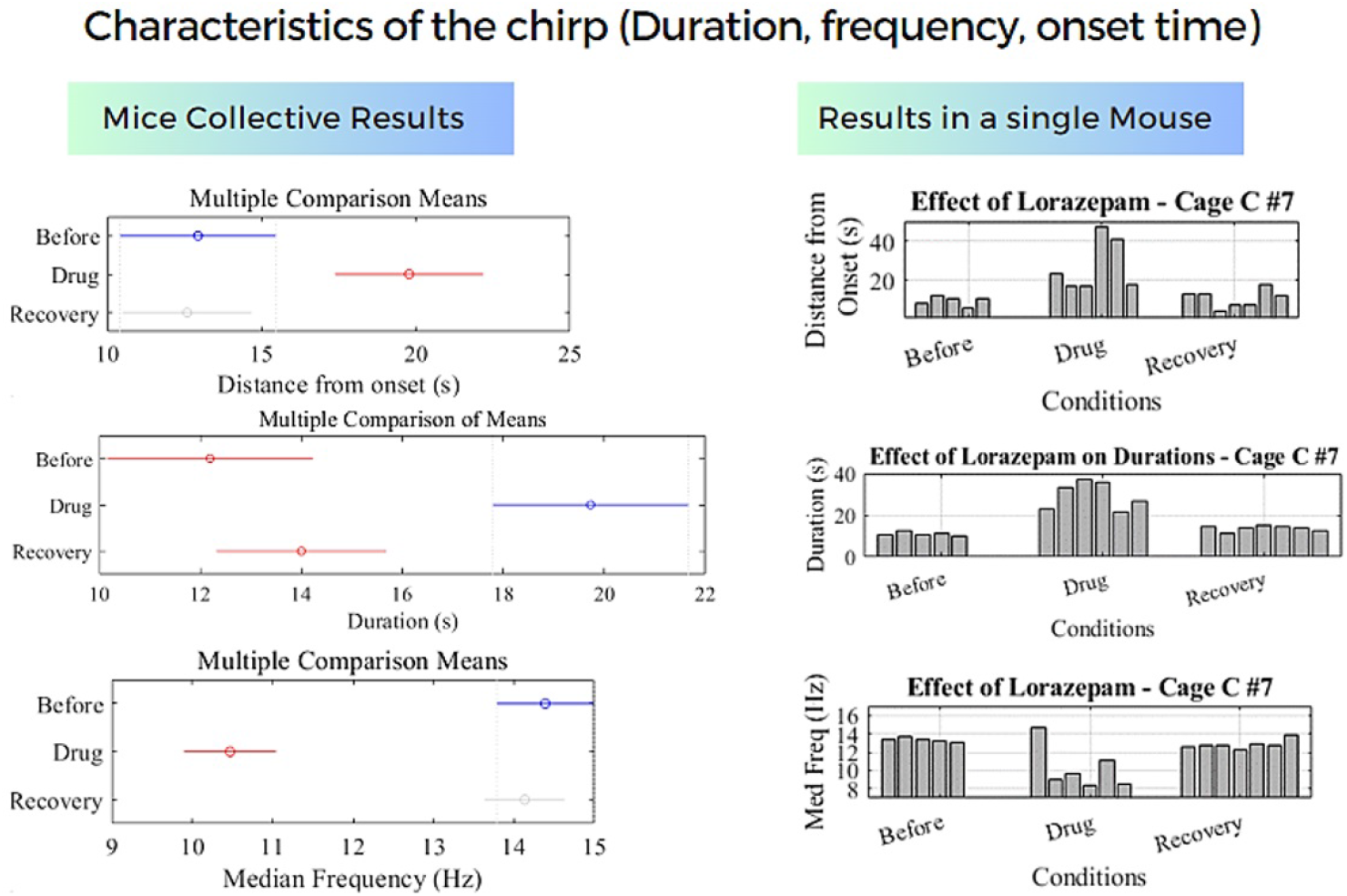
Chirp characteristics. Characteristics of the chirp before, during, and after lorazepam administration: On the left, the graphs display mean comparisons of chirp distance from ictal onset, chirp duration, and median chirp frequency, arranged respectively from top to bottom. These comparisons are made across all studied animals before, during, and after Lorazepam administration. On the right, the results for a single animal illustrate how Lorazepam leads to a slower and more stable chirp frequency, as well as a delay in the chirp’s occurrence during ictal activity.

## Discussion

For the first time, reliable characterization of chirp-like activities during ictal events has been investigated. It allowed us to characterize activities across various states of disease progression and drug application, with extracted features showing statistically significant differences among the different states. We reported notable changes in chirp features in response to drug application, indicating a pharmacological influence on this neurophysiological phenomenon. The key enabling factor behind this study is the availability of extensive, long-term recordings in epileptic mice. These recordings have allowed for the capture and analysis of hundreds to thousands of historical events. Unfortunately, such comprehensive data collection is not feasible in human recordings, where, for instance, only about three seizures can typically be obtained over several weeks in an epilepsy monitoring unit. Similarly, in previous animal studies, the documentation of large numbers of ictal events has been lacking, precluding thorough analysis. We anticipate that our findings will help understanding mechanisms underlying ictal-chirp and will collectively contribute to utilize chirps as both a diagnostic tool and a dynamic indicator of neurological responses to epilepsy severity and pharmacological interventions.

### Ictal-chirp patterns in other studies

Using a kindling method to induce seizures in rats, ***Sen et al. (2007***) observed a chirp pattern during ictal events using EEG recordings. It was observed that the frequency of chirps increased initially and decreased gradually as the seizure progressed. ***Sen et al. (2007***) reported that a kindling paradigm in rats can serve as an animal model of human temporal lobe epilepsy from the perspective of time-frequency variations.

A clinical study (***Li et al., 2020***) reported the presence of narrow-band gamma activity and narrow-band beta activity within a group of patients diagnosed with pharmaco-resistant focal epilepsy. This came through an evaluation conducted using Stereotactically implanted EEG (SEEG), an invasive pre-surgical procedure. Using sophisticated signal processing algorithms and visual inspection of time-frequency representation of SEEG signal during ictal events, ***Li et al. (2020***) showed that the beta-band pattern exhibits a tubular-like shape marked by a distinctive upward-downward trend. Another clinical study (***Gnatkovsky et al., 2011***) focused on presurgical evaluation of patients with drug-resistant focal epilepsy. The terms “correlation” and “coupling” between the power integral of very fast (220-270 Hz) and very slow (0.3-3 Hz) frequency band during ictal discharge were reported in one seizure type. Additionally, different frequency patterns in terms of frequency bands were observed in different seizure types. However, no information was provided regarding the time-frequency morphology of these patterns. The emphasis solely rested on the variations in power integral of frequency bands over time.

In another clinical study involving patients with mesial temporal lobe epilepsy, a distinctive signature event displaying a frequency evolution was observed prior to the onset of seizures (***Nieder-hauser et al., 2003***). These events were primarily situated within the 20-40 Hz frequency range. Despite the consistency in the shape of signature events before each seizure across patients, there were slight variations in offset frequencies and durations between individual seizures. The signature events typically had a duration exceeding 5 seconds. To identify these signature events within the EEG data, a process called temporal differentiation was applied. A mathematical function called the two-piece sign function was then applied on these differentiated values. This function assigned values of 1 or -1 to the differentiated data points based on whether they are positive or negative. Following this, a mathematical operation called the periodogram was applied, which involves using a sliding window and a fast Fourier transform to analyze the data in both time and frequency domains. This operation helps in capturing the energy distribution of the data over different frequency components. The resulting output was an energy spectrum that highlights significant events in the EEG data. This spectrum was normalized to have a consistent scale (***Nieder-hauser et al., 2003***). ***Schiff et al. (2000***) reported the presence of spectral chirps with decreasing frequency in intracranially recorded EEG from 6 patients with epileptic seizures. According to their results, the chirp intensity demonstrated a correlation with the spatial location of the seizure focus. The proximity of the electrodes to the point of seizure origination directly influenced the magnitude of the chirp intensity, with closer electrodes yielding stronger chirps. Based on their findings, chirps seemed to be effective in identifying seizures originating from both the neocortex and the hippocampus. The chirps in this study had a harmonic structure. In the realms of sonar and radar sensing, both sound and electromagnetic energy can traverse various paths from the source to the observer. These diverse pathways become more pronounced when the target is in motion. Essentially, phase delays emerge among signals of comparable frequencies, and the transmission’s numerous time delays give rise to distinct frequency bands – a phenomenon referred to as the ‘multipath’ effect. The smooth shifts within these bands, known as ‘chirps’ are directly connected to the object’s movement. Within neuronal structures like the cortex, a cluster of neurons might produce collective activity in an adjacent group, influenced by inherent delays in synaptic activation, involving both excitatory and inhibitory processes. Furthermore, the paths through which neuronal activity travels to reach the receiving electrode remain relatively stable yet vary in length. This study hypothesized that the migration of seizures as they propagate through the cortex could induce similar phenomena. The power spectra and spectrogram were used in this work to study the chirps. Chirp pattern in spectrogram was also used as a template to detect seizures (***Schiff et al., 2000***). In another clinical study based on EEG (***Feltane et al., 2013***), a chirp-like pattern was identified, appearing as a distinct ridge in the time-frequency representation. This ridge exhibited a linear decrease in frequency. The smoothed pseudo Wigner-Ville distribution was used in this study to generate the time-frequency representation. The Hough transform, a technique for detecting lines, circles, and ellipses within images, was employed to identify chirps within the time-frequency representation (***Feltane et al., 2013***). Stereo-EEG monitoring of patients with drug-resistant focal epilepsy in a clinical study revealed a down-chirping phenomenon (***Gnatkovsky et al., 2019a***). This involved a gradual decrease in high frequencies over a span of 5-10 seconds at the onset of an ictal event. A combination of computational and visual analyses was employed to detect this pattern. Additionally, the time-frequency representation was generated using a spectrogram. Chirps exhibiting a harmonic structure were also observed during the pre-seizure interval in the spectrogram of electrocorticogram (ECoG) data collected from the cortical surface of the brain (***Benedetto and Colella, 1995***).

### Ictal chirp as a biomarker for seizure onset zone detection

The primary goal of epilepsy surgery is to identify and remove the epileptogenic zone, the specific area in the brain where seizures originate. The resection volume, or the amount of brain tissue removed during surgery, remains an open question in epilepsy surgery. There is no universally accepted standard for resection volume estimation, nor is there enough scientifically defendable evidence to guide its optimal extent. Consequently, there is a significant need for a reliable indicator or biomarker to accurately determine the resection volume. The critical question is how to accurately estimate the resection margins with a minimum number of false positive (epileptogenic tissue outside the resection (the tissue was not removed during surgery)) and false negative channels (potentially epileptogenic tissue located inside the resection). One potential solution being explored is whether ictal-chirp can help identify the boundaries of the epileptogenic zone. Ictal-chirp has been reported to improve the accuracy of epilepsy surgery in humans (***Gnatkovsky et al., 2019b***; ***Grinenko et al., 2017***; ***Di Giacomo et al., 2024***). These studies provided evidence that chirp characteristics can differentiate the epileptogenic zone from areas of propagation.

### Limitations and Future work

Previous clinical and experimental studies observed a chirp-like pattern in seizure data, but they disagreed on when and how these chirps happened, partly because there weren’t enough seizure recordings to analyze. Our work here took advantage of a large dataset (epilepsy kindling model) and in-depth analyses to achieve a reliable assessment of chirp characteristics over time and different states. In future work, we can consider other types of epilepsy models besides kindling as well as seizures occurring in other neurological disease states. Further, one can consider using E-I network models to obtain mechanistic insights into the chirp phenomenon and its frequency variations by taking advantage of observed changes such as GABA_A_ enhancing drugs like lorazepam.

## Methods

### Experimental data description

Datasets of continuous EEG monitoring of mice undergoing extended kindling have been obtained and examined in previous studies (***Bin et al., 2017***; ***Liu et al., 2021***), including the effects of antiepileptic drugs (***Song et al., 2018***). We use several of these datasets and carry out the detailed data analyses of seizure events presented in this paper.

A description of the experimental protocol is provided here. C57 black mice (C57BL/6N, male) from Charles River Laboratory (Quebec, Canada), were housed under standard conditions (22-23 degrees celsius, 12-hour light on/off cycle starting at six AM). All experiments were approved by the University Health Network Animal Care Committee in line with Canadian Council on Animal Care guidelines. Each mouse was implanted with two pairs of twisted-wire bipolar electrodes. One pair of electrodes was positioned to the hippocampal CA3 region for kindling stimulation and local recordings, and another pair positioned to an ipsilateral or contralateral site. The latter included the contralateral hippocampal CA3, parietal cortex, piriform cortex and entorhinal cortex. A reference electrode was positioned to a frontal area. A train of stimuli at 60 Hz for two seconds was used for hippocampal kindling. Kindling stimuli were applied twice daily and ≥ 5 hours apart. Each stimulation episode lasted for a few minutes while the mouse was placed in a glass container for EEG-video monitoring. Control mice experienced twice daily handlings for 60 days. Local differential recordings through the twisted-wire bipolar electrodes were used to monitor evoked responses and spontaneous EEG activities. Evoked and spontaneous EEG signals were collected using two-channel or one-channel microelectrode AC amplifiers with extended head-stages (model 1800 or 3000, AM Systems; Sequim, Washington, USA). These amplifiers were set with an input frequency band of 0.1-1,000 Hz and an amplification gain of 1,000. A built-in notch filter at 60±3 Hz were used in some experiments. Amplifier output signals were digitized at 5,000 Hz (Digidata 1440A or 1550, Molecular Devices; Sunnyvale, California, USA). Individual mice underwent EEG-video monitoring for 24 hours after about 80, 100, 120 and 140 kindling stimulations. If ≥2 spontaneous recurrent seizure (SRS) events were observed in the 24-hour monitoring, no further kindling stimulation was applied, and EEG-video continued for up 6 days to assess SRS events in the early phase post kindling. EEG-video monitoring up to 6 consecutive days was resumed in some mice 6-12 weeks later to assess SRS in the late phase of post kindling. Evoked afterdepolarizations (ADs) and spontaneous discharges were recognized by repetitive spikes with simple and/or complex waveforms, amplitudes approximately 2 times the background signals and durations of ≥10 seconds. Most discharges began with low voltage fast (LVF) signals, which were followed by incremental rhythmic spikes and then sustained large-amplitude spikes with simple or complex waveforms. Discharge termination in most cases featured a sudden cessation of spike activity and a subsequent component of signal suppression lasting several seconds.

EEG recordings from nine mice with early and late evoked states were used for analysis in this study. Nine mice with spontaneous recurrent seizures were analyzed and six mice in which an antiepileptic drug (lorazepam) had been applied were also analyzed.

### Details of chirp detection method and analyses

Methods used for data analysis include time-frequency ridge extraction, chirp identification, and statistical testing. For time-frequency representation extraction, spectrogram calculation involved segmenting the signal, applying windowing functions, and performing the Discrete Fourier Transform to reveal the signal’s spectral evolution over time. Time-frequency ridges, representing dominant frequency components, were extracted using a forward-backward greedy algorithm with customizable parameters to penalize frequency changes and determine the number of ridges. Considering extracted ridges, the onset and offset of chirp was determined by analyzing power ratio of specific frequency bands and identifying chirp event based on a threshold.

#### Automated determination of onset and offset times for ictal-related chirps

Onset and offset times of ictal discharge-related chirps involves the following steps: 01) time-frequency analysis using spectrogram calculation; 02) Computing a smoothed power ratio trajectory of pre-defined high to low frequency bands over time, wherein high-frequency range is set between 10 Hz and 22 Hz, and low-frequency range is set between 1 Hz and 10 Hz (Frequency band of (1–10 Hz) was categorized as low frequency, while (8–20 Hz) was classified as high frequency. To prevent overlap, a distinction was made between <10 and >10 in the low and high-frequency bands (***Tort et al., 2018***)); 03) Applying a threshold of 1.5 to smoothed power ratio trajectory (The threshold was adjusted based on trial and error) ; 04) Identifying instances where the power ratio trajectory exceeds the threshold; 05) Confirming the significance of identified event through statistical analysis. The smoothed power ratio trajectory was segmented into two distinct categories: values surpassing the threshold and values equal to or below the threshold. By employing a two-sample t-test, a comparison of the means for these two groups was conducted. A very low p-value (p < 0.02) in the test results would indicate a significant difference between the groups. 06) Defining chirp onset as the moment when the power ratio trajectory significantly exceeds threshold and defining chirp offset as the moment when the power ratio trajectory significantly falls below threshold. The pre-defined range of high and low frequency bands may need fine-tuning through visual assessment because the range of low frequency and high frequency may vary slightly on a case-by-case basis. In FIGURE 8, the six stages involved in the automated detection of chirp onset and offset times are illustrated.

**Figure 8.**
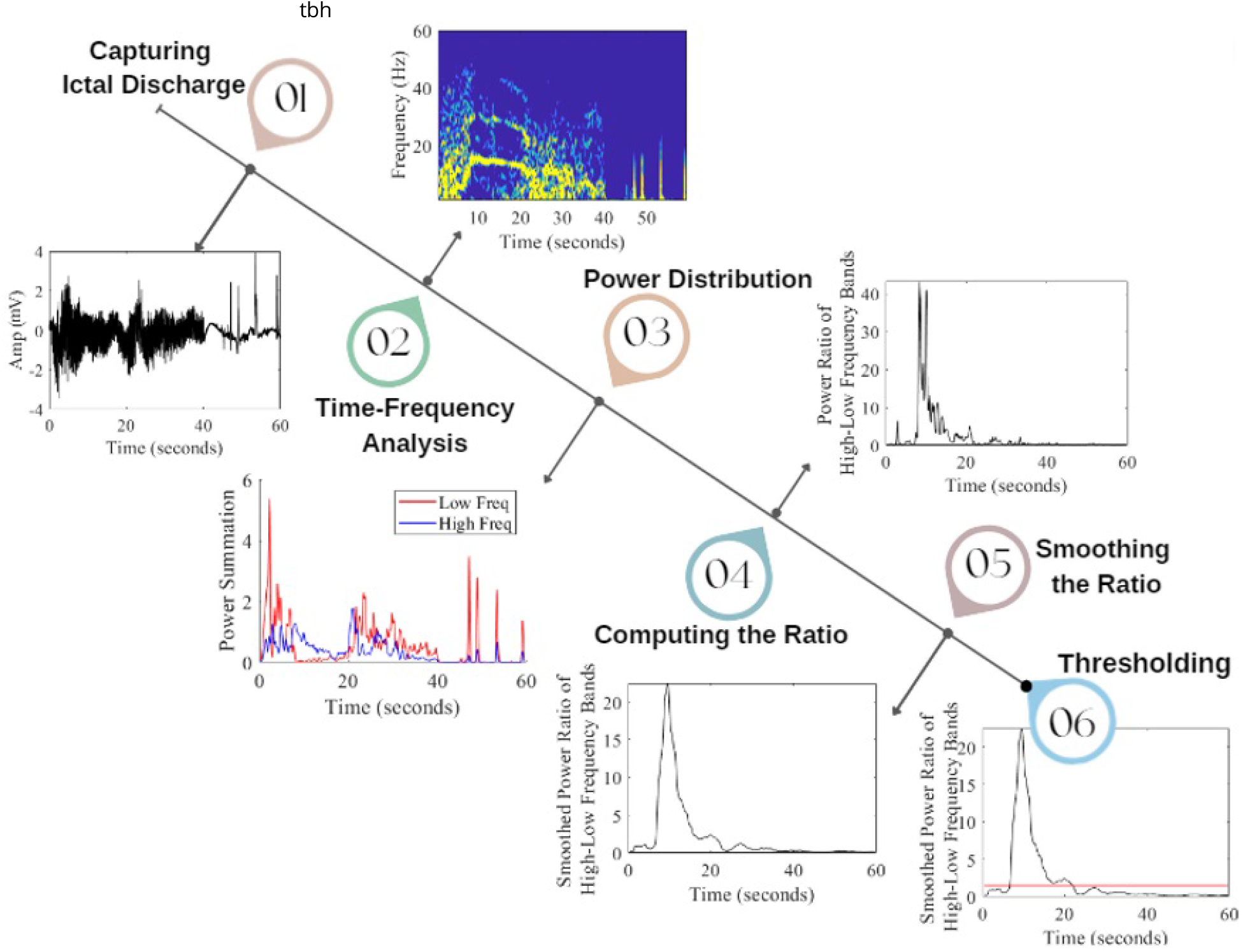
The six steps in automatic detection of chirp onset and offset times involve capturing ictal discharge (01), conducting time-frequency analysis (02), calculating power distribution in pre-defined bands (03), determining the high-to low-frequency power ratio (04), applying smoothing to the ratio trajectory (05), and setting thresholds (06). When evaluating the smoothed power ratio of high-low frequency bands over time, a threshold of 1.5 (red line), the high-frequency range being 10 Hz to 22 Hz and the low-frequency range spanning from 1 Hz to 10 Hz were considered. The smoothed power ratio trajectory was divided into two distinct groups: values exceeding the threshold and values equal to or below the threshold. Utilizing a two-sample t-test, a comparison of the means of these two groups revealed a very low *p*-value < 0.00001, indicating an extremely significant difference between the groups. The chirp’s initiation and termination were then defined as instances when the smoothed power ratio exceeded the specified threshold.

#### Automatic extraction of chirp morphology

After determining the onset and offset times, chirp morphologies were extracted by pulling out the dominant frequency ridge curve from the spectrogram (time-frequency representation) using a penalized forward-backward greedy algorithm between the determined onset and offset times of ictal-related chirp (***Bahador et al., 2021***). We performed an automated characterization of chirps by analyzing features of: duration of chirp, median frequency of chirp, and chirp onset (distance from ictal discharge onset).

#### Statistical tests

The statistical analysis investigated changes in chirp characteristics across various states of epilepsy by calculating mean values and confidence intervals. Significant differences between these means were assessed through pairwise comparisons using a multiple comparison test. The z-scores were computed to quantify the differences: a z-score greater than 2.58 indicates strong evidence against the null hypothesis (p < 0.01), a z-score above 1.96 suggests moderate evidence (p < 0.05), and a z-score exceeding 1.64 indicates some evidence (p < 0.10); results below these thresholds were considered not significant (n.s.). Additionally, a series of two-sample t-tests assuming unequal variances were performed to compare chirp durations between specific pairs of states, including early evoked vs. late evoked, late evoked vs. spontaneous recovery state (SRS), and various comparisons involving the drug state and recovery, thereby elucidating how chirp characteristics change as epilepsy progresses or responds to treatment.

## Supplementary Tables

**Table 1.** t-Test: Two-Sample Assuming Unequal Variances (Chirp Duration, Early Evoked vs. Late Evoked)

|  | Early Evoked | Late Evoked |
| --- | --- | --- |
| Mean | 2.354166667 | 4.135555556 |
| Variance | 1.305199275 | 3.00779798 |
| Observations | 24 | 45 |
| Hypothesized Mean Difference | 0 | 0 |
| df | 64 | 64 |
| t Stat | -5.116414991 | -5.116414991 |
| P(T≤t) one-tail | 1.52913E-06 | 1.52913E-06 |
| t Critical one-tail | 1.669013025 | 1.669013025 |
| P(T≤t) two-tail | 3.05825E-06 | 3.05825E-06 |
| t Critical two-tail | 1.997729654 | 1.997729654 |

**Table 2.** t-Test: Two-Sample Assuming Unequal Variances (Chirp Duration, Late Evoked vs. SRS)

|  | Late Evoked | SRS |
| --- | --- | --- |
| Mean | 4.135555556 | 9.613636364 |
| Variance | 3.00779798 | 9.520272727 |
| Observations | 45 | 66 |
| Hypothesized Mean Difference | 0 | 0 |
| df | 106 | 106 |
| t Stat | -11.92334727 | -11.92334727 |
| P(T≤t) one-tail | 1.33385E-21 | 1.33385E-21 |
| t Critical one-tail | 1.659356034 | 1.659356034 |
| P(T≤t) two-tail | 2.6677E-21 | 2.6677E-21 |
| t Critical two-tail | 1.982597262 | 1.982597262 |

**Table 3.** t-Test: Two-Sample Assuming Unequal Variances (Chirp Duration, SRS vs. Late SRS)

|  | SRS | Late SRS |
| --- | --- | --- |
| Mean | 9.613636364 | 13.35882353 |
| Variance | 9.520272727 | 5.365073529 |
| Observations | 66 | 17 |
| Hypothesized Mean Difference | 0 | 0 |
| df | 32 | 32 |
| t Stat | -5.522943116 | -5.522943116 |
| P(T≤t) one-tail | 2.16857E-06 | 2.16857E-06 |
| t Critical one-tail | 1.693888748 | 1.693888748 |
| P(T≤t) two-tail | 4.33714E-06 | 4.33714E-06 |
| t Critical two-tail | 2.036933343 | 2.036933343 |

**Table 4.**
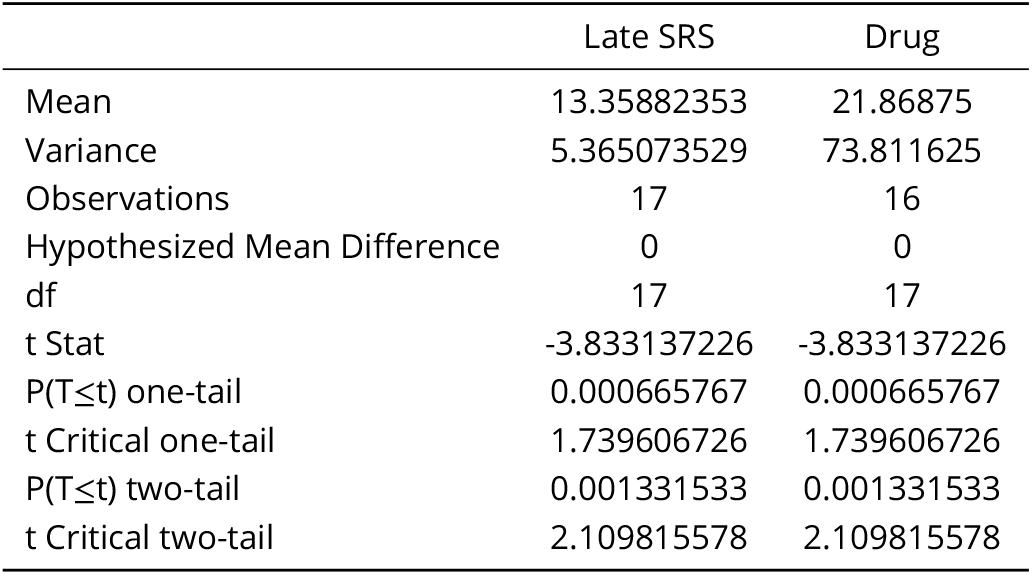
t-Test: Two-Sample Assuming Unequal Variances (Chirp Duration, Late SRS vs. Drug)

**Table 5.**
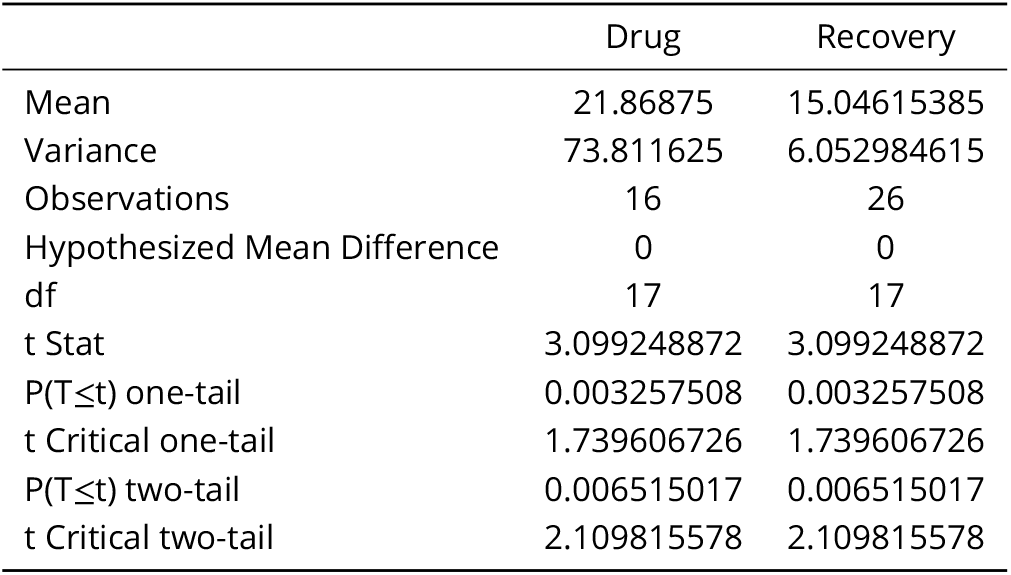
t-Test: Two-Sample Assuming Unequal Variances (Chirp Duration, Drug vs. Recovery)

**Table 6.** t-Test: Two-Sample Assuming Unequal Variances (Chirp Duration, SRS vs. Recovery)

|  | SRS | Recovery |
| --- | --- | --- |
| Mean | 9.613636364 | 15.04615385 |
| Variance | 9.520272727 | 6.052984615 |
| Observations | 66 | 26 |
| Hypothesized Mean Difference | 0 | 0 |
| df | 57 | 57 |
| t Stat | -8.847071841 | -8.847071841 |
| P(T≤t) one-tail | 1.37337E-12 | 1.37337E-12 |
| t Critical one-tail | 1.672028888 | 1.672028888 |
| P(T≤t) two-tail | 2.74675E-12 | 2.74675E-12 |
| t Critical two-tail | 2.002465459 | 2.002465459 |

**Table 7.** t-Test: Two-Sample Assuming Unequal Variances (Chirp Duration, Late SRS vs. Recovery)

|  | Late SRS | Recovery |
| --- | --- | --- |
| Mean | 13.35882353 | 15.04615385 |
| Variance | 5.365073529 | 6.052984615 |
| Observations | 17 | 26 |
| Hypothesized Mean Difference | 0 | 0 |
| df | 36 | 36 |
| t Stat | -2.278513053 | -2.278513053 |
| P(T≤t) one-tail | 0.014365616 | 0.014365616 |
| t Critical one-tail | 1.688297714 | 1.688297714 |
| P(T≤t) two-tail | 0.028731232 | 0.028731232 |
| t Critical two-tail | 2.028094001 | 2.028094001 |

**Table 8.** t-Test: Two-Sample Assuming Unequal Variances (Pre-Drug State vs Drug State)

|  | Duration Pre-Drug State (s) | Duration Drug State (s) |
| --- | --- | --- |
| Mean | 12.1952381 | 19.72608696 |
| Variance | 11.05447619 | 73.38928854 |
| Observations | 21 | 23 |
| Hypothesized Mean Difference | 0 | 0 |
| df | 29 | 29 |
| t Stat | -3.906011331 | -3.906011331 |
| P(T≤t) one-tail | 0.000258035 | 0.000258035 |
| t Critical one-tail | 1.699127027 | 1.699127027 |
| P(T≤t) two-tail | 0.00051607 | 0.00051607 |
| t Critical two-tail | 2.045229642 | 2.045229642 |

**Table 9.** t-Test: Two-Sample Assuming Unequal Variances (Drug State vs Recovery State)

|  | Duration Drug State (s) | Duration Recovery State (s) |
| --- | --- | --- |
| Mean | 19.72608696 | 14.00322581 |
| Variance | 73.38928854 | 11.25498925 |
| Observations | 23 | 31 |
| Hypothesized Mean Difference | 0 | 0 |
| df | 27 | 27 |
| t Stat | 3.035711083 | 3.035711083 |
| P(T≤t) one-tail | 0.002631562 | 0.002631562 |
| t Critical one-tail | 1.703288446 | 1.703288446 |
| P(T≤t) two-tail | 0.005263124 | 0.005263124 |
| t Critical two-tail | 2.051830516 | 2.051830516 |

**Table 10.** t-Test: Two-Sample Assuming Unequal Variances (Pre-Drug State vs Recovery State)

|  | Duration Pre-Drug State (s) | Duration Recovery State (s) |
| --- | --- | --- |
| Mean | 12.1952381 | 14.00322581 |
| Variance | 11.05447619 | 11.25498925 |
| Observations | 21 | 31 |
| Hypothesized Mean Difference | 0 | 0 |
| df | 43 | 43 |
| t Stat | -1.917036402 | -1.917036402 |
| P(T≤t) one-tail | 0.030946631 | 0.030946631 |
| t Critical one-tail | 1.681070703 | 1.681070703 |
| P(T≤t) two-tail | 0.061893262 | 0.061893262 |
| t Critical two-tail | 2.016692199 | 2.016692199 |

## Acknowledgments

Additional information can be given in the template, such as to not include funder information in the acknowledgments section.

---

**Figure 1—figure supplement 1.**
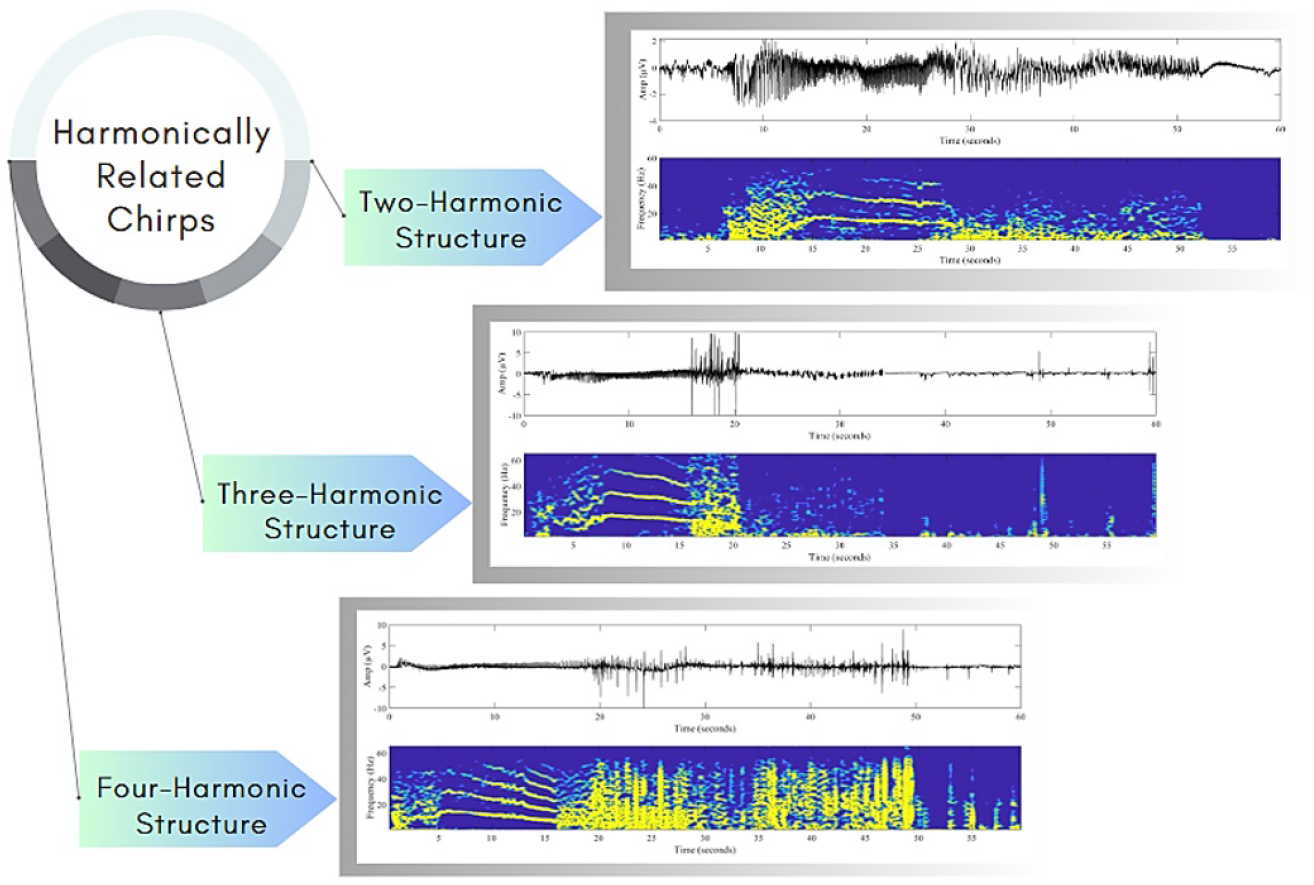
Chirps exhibiting harmonic relationships. Harmonically related chirps may reflect the resonant properties of the mechanism that generates them.

**Figure 1—figure supplement 2.**
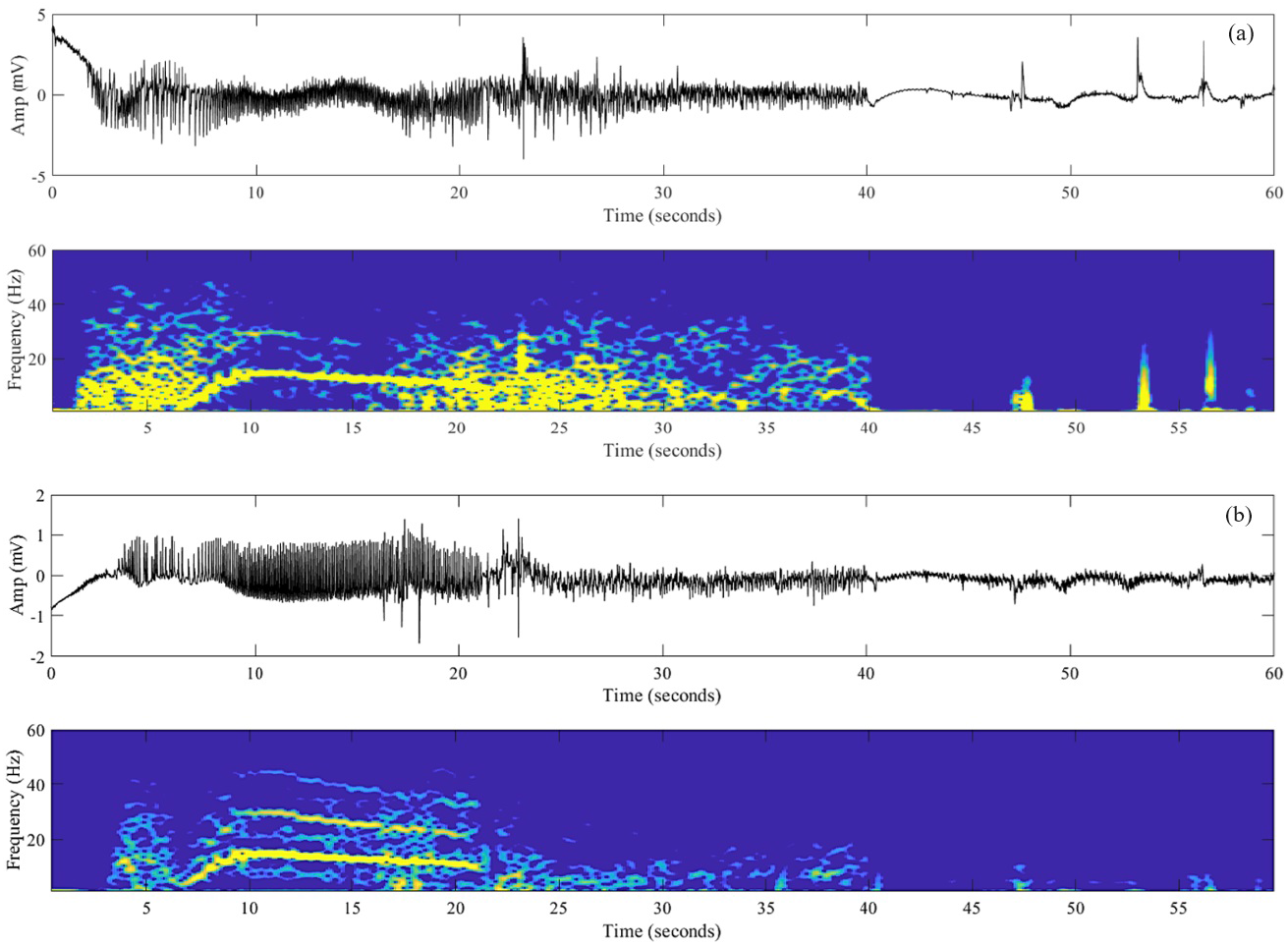
Chirp patterns in two different brain regions. Concurrent chirp patterns across distinct brain regions during spontaneous recurrent seizure in the same animal. (a) Ictal discharge of hippocampus. (b) Ictal discharge of piriform cortex.

**Figure 1—figure supplement 3.**
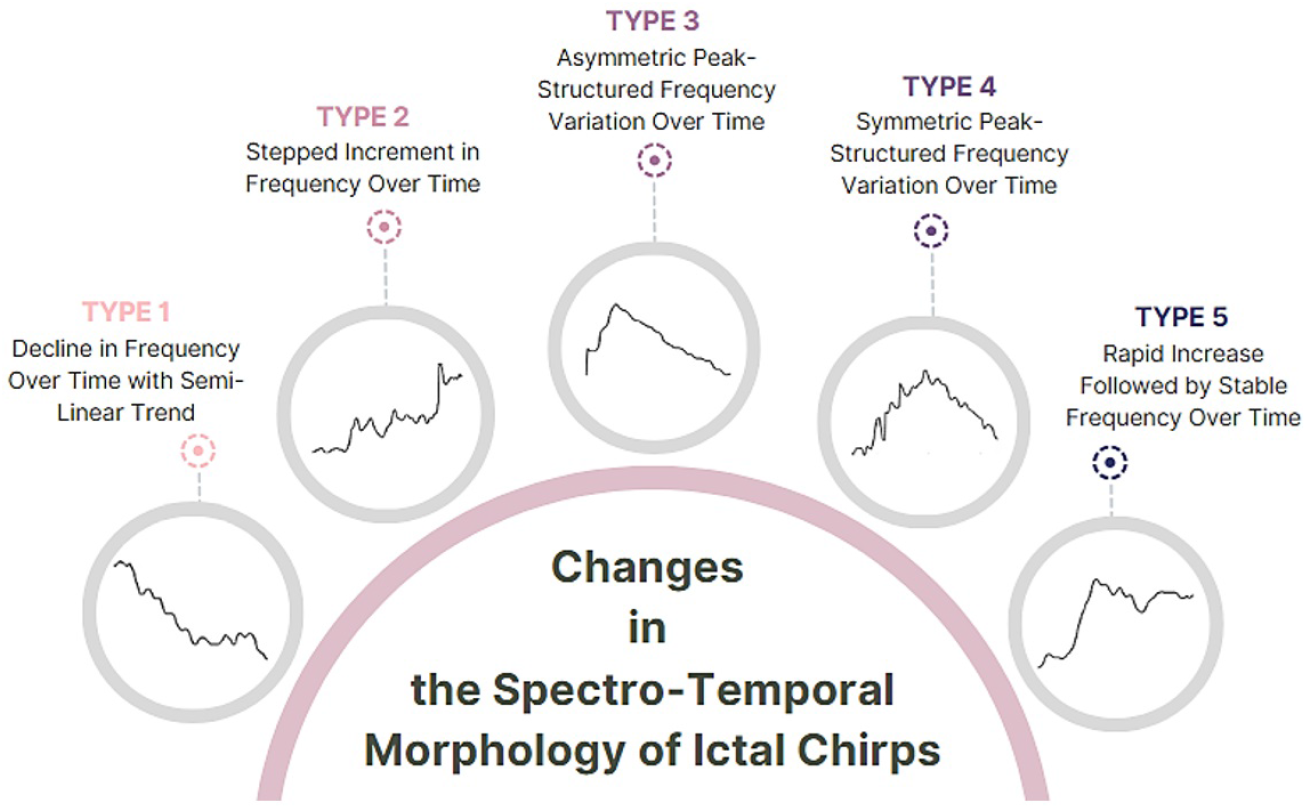
Spectro-temporal morphology of ictal chirps. Various patterns in the spectro-temporal morphology of ictal chirps, including declining frequency with a semi-linear trend, stepped frequency increments, asymmetric and symmetric peak-structured frequency variations over time, and rapid initial increases followed by stable frequencies. They can be categorized into five distinct types based on the evolution of their frequency characteristics over time. Type 1 exhibits a decline in frequency with a semi-linear trend; Type 2 displays a stepped increment in frequency over time; Type 3 features asymmetric peak-structured frequency variations; Type 4 showcases symmetric peak-structured frequency changes; Type 5 is characterized by a rapid initial increase in frequency followed by a stable frequency profile over time. These patterns are the ones most frequently observed in our dataset, as determined by visual assessments.

**Figure 3—figure supplement 1.**
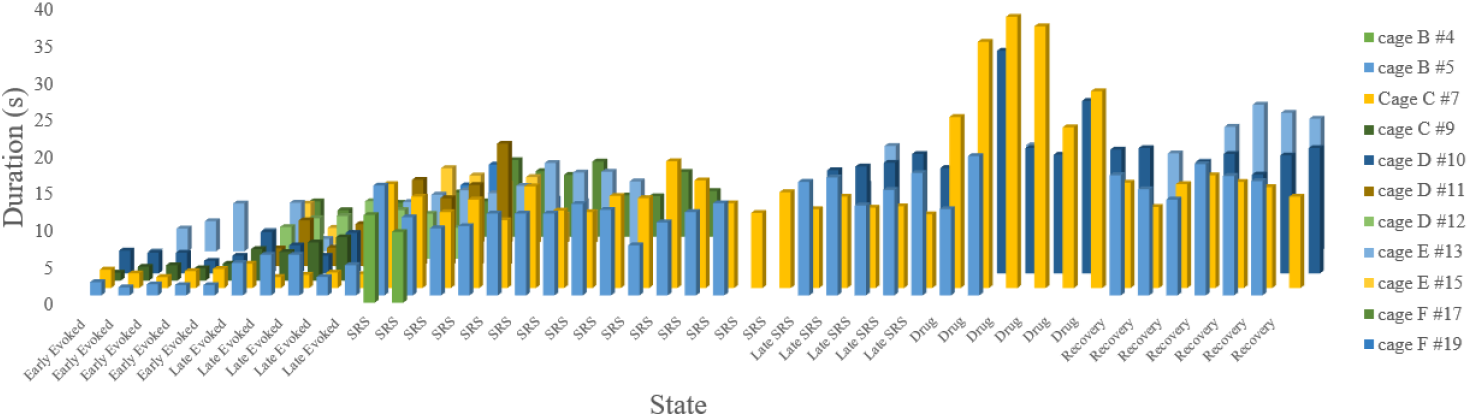
Variation in chirp duration (s) across different animals over time: Early Evoked, Late Evoked, SRS, Late SRS, Drug, and Recovery.

**Figure 3—figure supplement 2.**
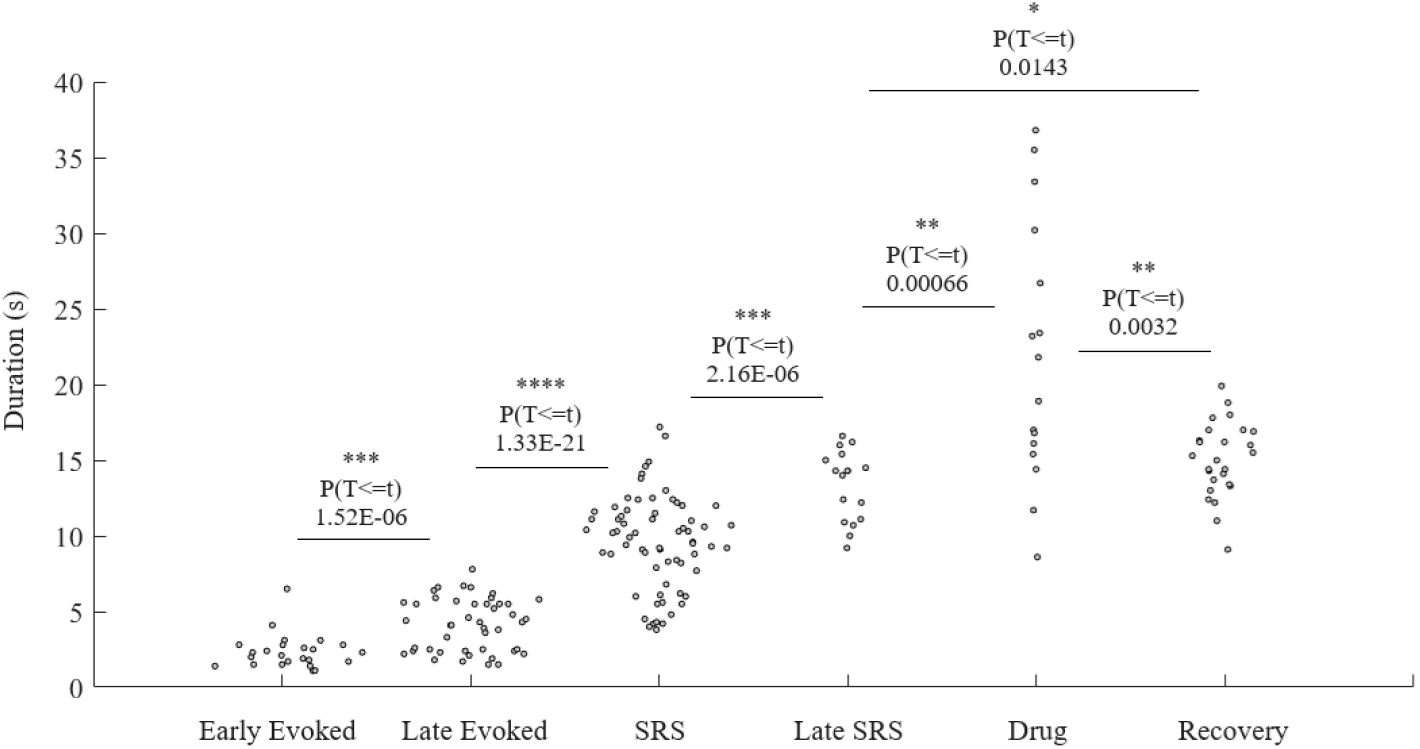
Statistical difference in chirp duration (s) across different states: Early Evoked, Late Evoked, SRS, Late SRS, Drug, and Recovery.

**Figure 3—figure supplement 3.**
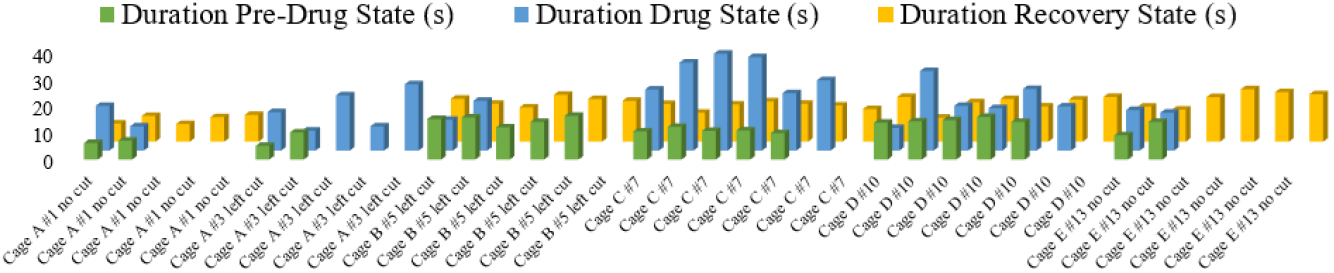
Chirp duration of Pre-Drug, Drug, and Recovery states in different animals.

**Figure 4—figure supplement 1.**
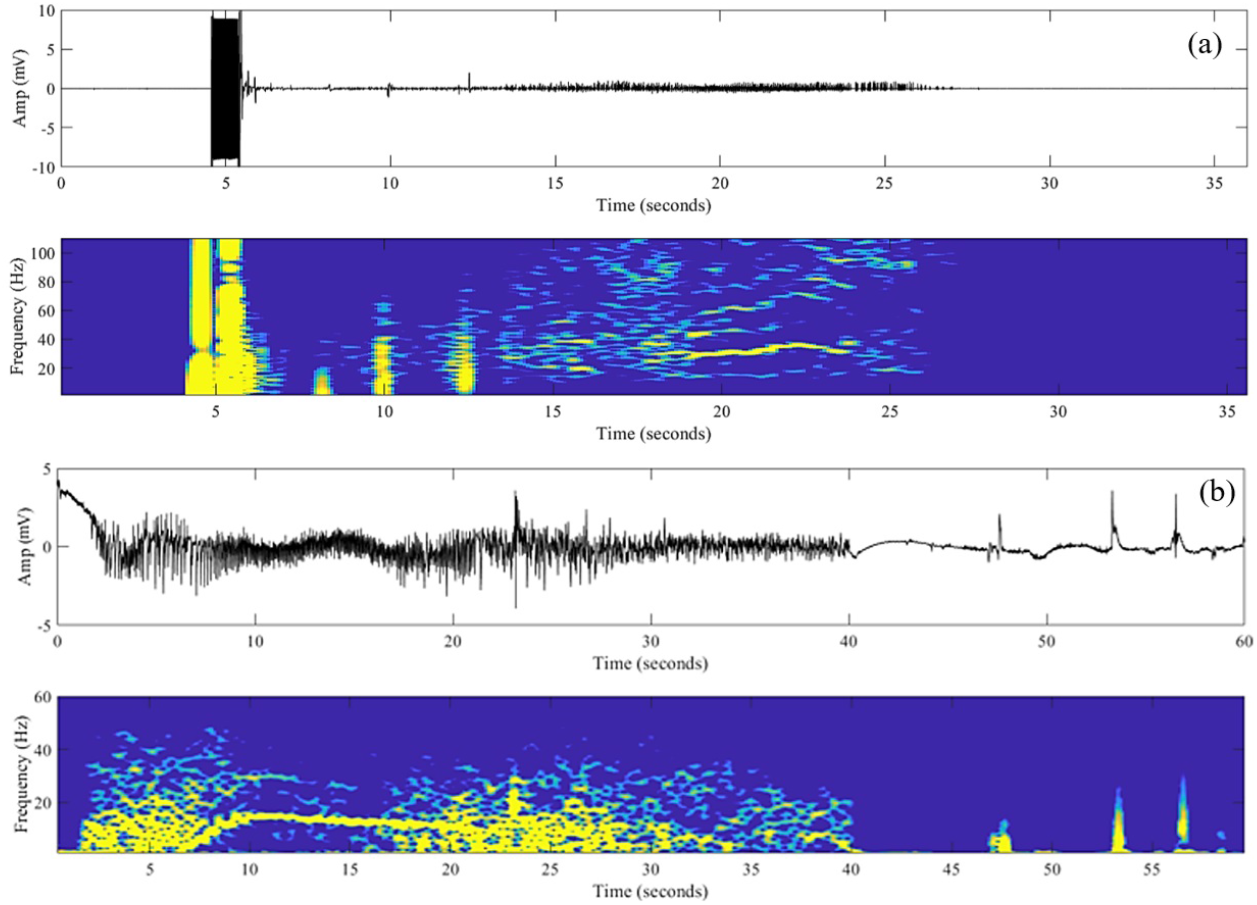
Chirp occurrence differences. Temporal relationship between chirp occurrence and ictal termination in an early stage evoked discharge (a) compared to a later SRS one (b).

## Notes

### Competing Interest Statement

The authors have declared no competing interest.

## References

Bahador, N., Jokelainen, J., Mustola, S., and Kortelainen, J. (2021). Multimodal spatio-temporal-spectral fusion for deep learning applications in physiological time series processing: A case study in monitoring the depth of anesthesia. Information Fusion, 73:125–143.

Benedetto, J. J. and Colella, D. (1995). Wavelet analysis of spectrogram seizure chirps. In Wavelet Applications in Signal and Image Processing III, volume 2569, pages 512–521. SPIE.

Bin, N.-R., Song, H., Wu, C., Lau, M., Sugita, S., Eubanks, J. H., and Zhang, L. (2017). Continuous monitoring via tethered electroencephalography of spontaneous recurrent seizures in mice. Frontiers in behavioral neuroscience, 11:172.

Brunel, N. (2000). Dynamics of Sparsely Connected Networks of Excitatory and Inhibitory Spiking Neurons. Journal of Computational Neuroscience, 8(3):183–208.

Brunel, N. and Wang, X.-J. (2003). What Determines the Frequency of Fast Network Oscillations With Irregular Neural Discharges? I. Synaptic Dynamics and Excitation-Inhibition Balance. Journal of Neurophysiology, 90(1):415–430.

Chen, X., van Gerven, J., Cohen, A., and Jacobs, G. (2019). Human pharmacology of positive gaba-a subtype-selective receptor modulators for the treatment of anxiety. Acta Pharmacologica Sinica, 40(5):571–582.

Coan, A. C. and Cendes, F. (2013). Epilepsy as progressive disorders: what is the evidence that can guide our clinical decisions and how can neuroimaging help? Epilepsy & Behavior, 26(3):313–321.

de Curtis, M. and Avoli, M. (2016). GABAergic networks jump-start focal seizures. Epilepsia, 57(5):679–687.

Di Giacomo, R., Burini, A., Chiarello, D., Pelliccia, V., Deleo, F., Garbelli, R., de Curtis, M., Tassi, L., and Gnatkovsky, V. (2024). Ictal fast activity chirps as markers of the epileptogenic zone. Epilepsia, 65(6).

El-Hayek, Y. H., Wu, C., Chen, R., Al-Sharif, A. R., Huang, S., Patel, N., Du, C., Ruff, C. A., Fehlings, M. G., Carlen, P. L., et al. (2011). Acute postischemic seizures are associated with increased mortality and brain damage in adult mice. Cerebral Cortex, 21(12):2863–2875.

Feltane, A., Bartels, G. B., Boudria, Y., and Besio, W. (2013). Analyzing the presence of chirp signals in the electroencephalogram during seizure using the reassignment time-frequency representation and the hough transform. In 2013 6th International IEEE/EMBS Conference on Neural Engineering (NER), pages 186–189. IEEE.

Freeman, J. M., Vining, E. P., and J., P. D. (1993). Seizures and epilepsy in childhood : a guide for parents.

Frosz, M. H. and Andersen, P. E. (2007). Can pulse broadening be stopped? Nature Photonics, 1(11):611–612.

Gnatkovsky, V., Francione, S., Cardinale, F., Mai, R., Tassi, L., Lo Russo, G., and De Curtis, M. (2011). Identification of reproducible ictal patterns based on quantified frequency analysis of intracranial eeg signals. Epilepsia, 52(3):477–488.

Gnatkovsky, V., Pelliccia, V., de Curtis, M., and Tassi, L. (2019a). Two main focal seizure patterns revealed by intracerebral electroencephalographic biomarker analysis. Epilepsia, 60(1):96–106.

Gnatkovsky, V., Pelliccia, V., de Curtis, M., and Tassi, L. (2019b). Two main focal seizure patterns revealed by intracerebral electroencephalographic biomarker analysis. Epilepsia, 60(1):96–106.

Grinenko, O., Li, J., Mosher, J. C., Wang, I. Z., Bulacio, J. C., Gonzalez-Martinez, J., Nair, D., Najm, I., Leahy, R. M., and Chauvel, P. (2017). A fingerprint of the epileptogenic zone in human epilepsies. Brain, 141(1):117–131.

Grinenko, O., Li, J., Mosher, J. C., Wang, I. Z., Bulacio, J. C., Gonzalez-Martinez, J., Nair, D., Najm, I., Leahy, R. M., and Chauvel, P. (2018). A fingerprint of the epileptogenic zone in human epilepsies. Brain, 141(1):117–131.

Jeffrey, M., Lang, M., Gane, J., Chow, E., Wu, C., and Zhang, L. (2014). Novel anticonvulsive effects of progesterone in a mouse model of hippocampal electrical kindling. Neuroscience, 257:65–75.

Kienitz, R., Kay, L., Beuchat, I., Gelhard, S., von Brauchitsch, S., Mann, C., Lucaciu, A., Schäfer, J.-H., Siebenbrodt, K., Zöllner, J.-P., et al. (2022). Benzodiazepines in the management of seizures and status epilepticus: A review of routes of delivery, pharmacokinetics, efficacy, and tolerability. CNS drugs, 36(9):951–975.

Kurbatova, P., Wendling, F., Kaminska, A., Rosati, A., Nabbout, R., Guerrini, R., Dulac, O., Pons, G., Cornu, C., Nony, P., et al. (2016). Dynamic changes of depolarizing gaba in a computational model of epileptogenic brain: Insight for dravet syndrome. Experimental neurology, 283:57–72.

Li, J., Grinenko, O., Mosher, J. C., Gonzalez-Martinez, J., Leahy, R. M., and Chauvel, P. (2020). Learning to define an electrical biomarker of the epileptogenic zone. Human Brain Mapping, 41(2):429–441.

Liu, H., Tufa, U., Zahra, A., Chow, J., Sivanenthiran, N., Cheng, C., Liu, Y., Cheung, P., Lim, S., Jin, Y., et al. (2021). Electrographic features of spontaneous recurrent seizures in a mouse model of extended hippocampal kindling. Cerebral Cortex Communications, 2(1):tgab004.

Miri, M. L., Vinck, M., Pant, R., and Cardin, J. A. (2018). Altered hippocampal interneuron activity precedes ictal onset. eLife, 7:e40750.

Niederhauser, J. J., Esteller, R., Echauz, J., Vachtsevanos, G., and Litt, B. (2003). Detection of seizure precursors from depth-eeg using a sign periodogram transform. IEEE Transactions on Biomedical Engineering, 50(4):449– 458.

Proix, T., Jirsa, V. K., Bartolomei, F., Guye, M., and Truccolo, W. (2018). Predicting the spatiotemporal diversity of seizure propagation and termination in human focal epilepsy. Nature communications, 9(1):1088.

Reiter, J. M. and Andrews, D. J. (2000). A neurobehavioral approach for treatment of complex partial epilepsy: efficacy. Seizure, 9(3):198–203.

Rich, S., Chameh, H. M., Rafiee, M., Ferguson, K., Skinner, F. K., and Valiante, T. A. (2020). Inhibitory Network Bistability Explains Increased Interneuronal Activity Prior to Seizure Onset. Frontiers in Neural Circuits, 13.

Sanna, E., Talani, G., Busonero, F., Pisu, M. G., Purdy, R. H., Serra, M., and Biggio, G. (2004). Brain Steroidogenesis Mediates Ethanol Modulation of GABAA Receptor Activity in Rat Hippocampus. Journal of Neuroscience, 24(29):6521–6530. Publisher: Society for Neuroscience Section: Cellular/Molecular.

Schiff, S. J., Colella, D., Jacyna, G. M., Hughes, E., Creekmore, J. W., Marshall, A., Bozek-Kuzmicki, M., Benke, G., Gaillard, W. D., Conry, J., et al. (2000). Brain chirps: spectrographic signatures of epileptic seizures. Clinical Neurophysiology, 111(6):953–958.

Sen, A., Kubek, M., and Shannon, H. (2007). Analysis of seizure eeg in kindled epileptic rats. Computational and Mathematical Methods in Medicine, 8(4):225–234.

Song, H., Tufa, U., Chow, J., Sivanenthiran, N., Cheng, C., Lim, S., Wu, C., Feng, J., Eubanks, J. H., and Zhang, L. (2018). Effects of antiepileptic drugs on spontaneous recurrent seizures in a novel model of extended hippocampal kindling in mice. Frontiers in Pharmacology, 9:451.

Stover, K. R., Lim, S., Zhou, T.-L., Stafford, P. M., Chow, J., Li, H., Sivanenthiran, N., Mylvaganam, S., Wu, C., Weaver, D. F., et al. (2017). Susceptibility to hippocampal kindling seizures is increased in aging c57 black mice. IBRO reports, 3:33–44.

Tort, A. B., Ponsel, S., Jessberger, J., Yanovsky, Y., Brankack, J., and Draguhn, A. (2018). Parallel detection of theta and respiration-coupled oscillations throughout the mouse brain. Scientific reports, 8(1):6432.

Trevorrow, T. (2006). Air travel and seizure frequency for individuals with epilepsy. Seizure, 15(5):320–327.

Truccolo, W., Ahmed, O. J., Harrison, M. T., Eskandar, E. N., Cosgrove, G. R., Madsen, J. R., Blum, A. S., Potter, N. S., Hochberg, L. R., and Cash, S. S. (2014). Neuronal Ensemble Synchrony during Human Focal Seizures. Journal of Neuroscience, 34(30):9927–9944. Publisher: Society for Neuroscience Section: Articles.

Truccolo, W., Donoghue, J. A., Hochberg, L. R., Eskandar, E. N., Madsen, J. R., Anderson, W. S., Brown, E. N., Halgren, E., and Cash, S. S. (2011). Single-neuron dynamics in human focal epilepsy. Nat Neurosci, advance online publication.

Wu, C., Wais, M., Sheppy, E., del Campo, M., and Zhang, L. (2008). A glue-based, screw-free method for implantation of intra-cranial electrodes in young mice. Journal of neuroscience methods, 171(1):126–131.

Y. Ho, E. C. and Truccolo, W. (2016). Interaction between synaptic inhibition and glial-potassium dynamics leads to diverse seizure transition modes in biophysical models of human focal seizures. Journal of Computational Neuroscience, 41(2):225–244.

